# Allergy-induced systemic inflammation impairs tendon quality

**DOI:** 10.1101/2021.07.02.450910

**Authors:** Christine Lehner, Gabriel Spitzer, Patrick Langthaler, Dominika Jakubecova, Barbara Klein, Nadja Weissenbacher, Andrea Wagner, Renate Gehwolf, Eugen Trinka, Bernhard Iglseder, Bernhard Paulweber, Ludwig Aigner, Sebastien Couillard-Després, Richard Weiss, Herbert Tempfer, Andreas Traweger

## Abstract

Treatment of tendinopathies still present a major challenge, since the aetiology of the disease remains poorly understood. To determine whether the systemic inflammation accompanying predisposing factors including rheumatoid arthritis, diabetes or smoking contributes to the onset of tendinopathy, we studied the effect of a systemic inflammation induced by an allergic episode on tendon properties. To this end, we elicited an allergic response in mice by exposing them to a plant allergen and subsequently analysed both their flexor and Achilles tendons. Biomechanical testing and histological analysis revealed that tendons from allergic mice not only showed a significant reduction of both elastic modulus and tensile stress, but also alterations of the tendon matrix. Moreover, 3D tendon-like constructs treated with sera from allergic mice displayed a matrix-remodelling expression profile and the expression of macrophage-associated markers and matrix metalloproteinase 2 (MMP2) was increased in allergic Achilles tendons. Analysing data from an epidemiologic study comprising data from more than 10.000 persons, we found that persons suffering from an allergic condition appeared to have an increased propensity to develop a tendinopathy.

**Graphical abstract:** 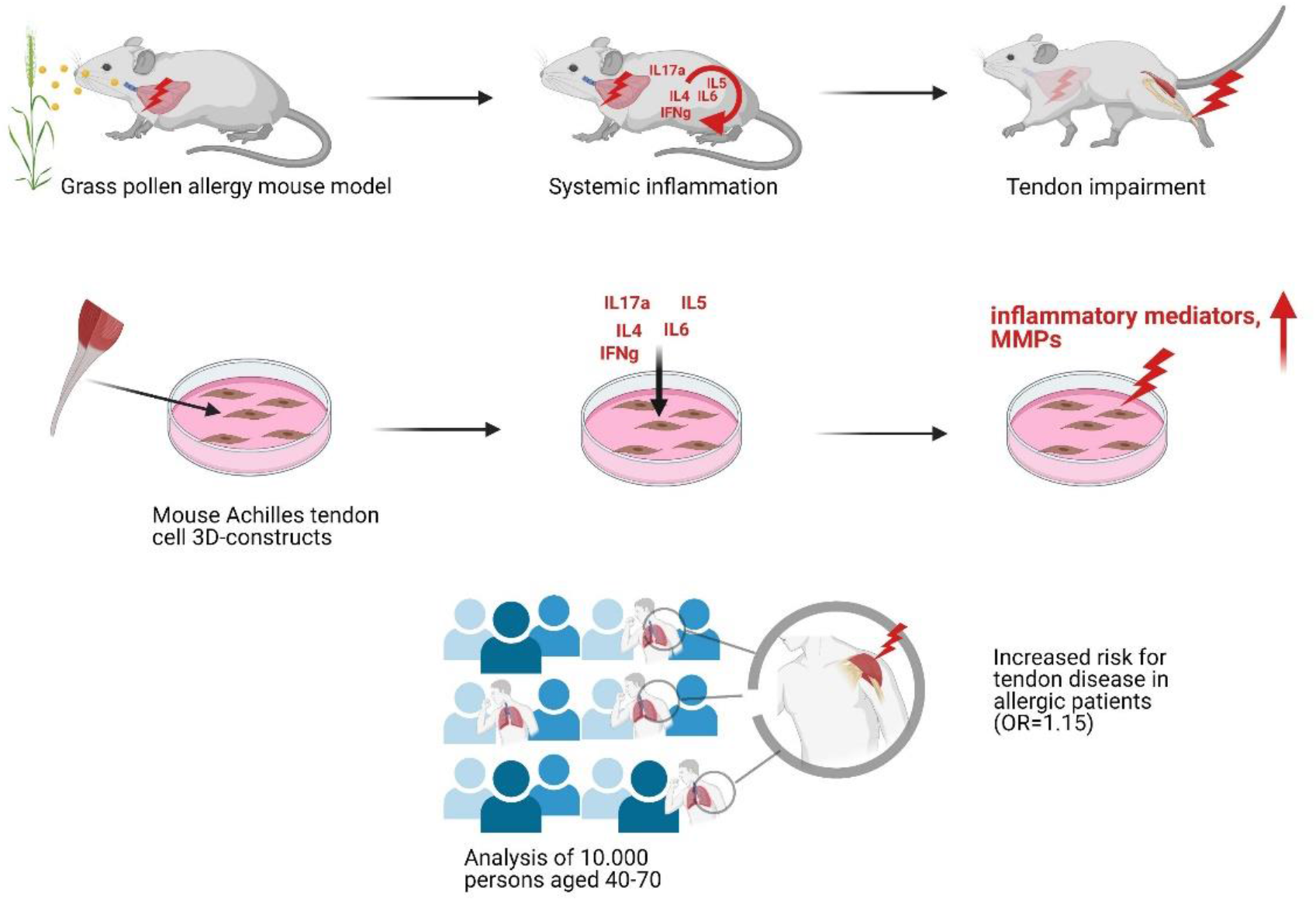

## Introduction

Tendon disorders are common and besides patients’ pain and disability entail considerable healthcare costs and loss of productivity [1]. Apart from laceration or acute overload-trauma in healthy tendons, tendon ruptures frequently occur in tendinopathic tendons, which undergo tissue degeneration due to overuse conditions or various intrinsic factors (e.g. age) [2]. Unfavourably, tendon tissue is known to possess a very poor healing capacity and in case of a rupture the repair tissue is of inferior quality displaying reduced mechanical properties with an increased risk in retears, the incidence of retears e.g. in rotator cuff tendons being more than 50 % [3, 4].

Tendinopathies are characterized by redness, swelling, pain and a limited range of motion. Typical features of tendinopathic tendons include fatty and mucoid infiltrations, calcific deposits, neurovascular ingrowth, hyperproliferation, and disoriented collagen fibers. The underlying cellular and molecular mechanisms are believed to be multifactorial. Yet, they are still poorly understood, but consensus exists about the fact that overuse, repetitive strain, and impingement are key drivers. Additionally, other extrinsic factors such as smoking, age, and the use of certain drugs such as corticosteroids or fluoroquinolones are considered further predisposing factors for tendon pathologies [5]. To what degree and at which stage inflammatory processes are involved in the development of tendinopathies is still a matter of debate, but over recent years increasing evidence has been pointing towards an inflammatory involvement [6–9].

Interestingly, many diseases including metabolic disorders such as diabetes, obesity, hypercholesterolaemia and hyperuricaemia have been described to be associated with tendinopathy [10–14]. Also, spontaneous quadriceps ruptures have been reported from patients suffering from primary and secondary hyperparathyroidism and chronic renal failure [15, 16]. The majority of these conditions commonly go along with a systemic, chronic low-grade inflammation [17], characterized by a two to three-fold increase in circulating cytokines and numerous other markers of immune system activity [18]. Consistently, tendon ruptures have been described to occur in a variety of autoimmune diseases including systemic lupus erythematosus, rheumatoid arthritis, gout and psoriasis, all of which are accompanied by a more or less pronounced chronic inflammation [19–21]. Interestingly, in a recent population-based cohort study significant associations between common allergic diseases and incident rheumatoid arthritis (RA) have been reported [22], indicating that allergic diseases and RA might share a similar underlying etiology related to chronic inflammatory responses.

Although allergies are linked to systemic subclinical inflammation, it has not been investigated so far whether persons suffering from allergies e.g. allergic rhinitis or asthma also have a higher risk to develop tendinopathies [23]. Since patients suffering under these allergic conditions frequently receive corticosteroid medication (often over long time periods) and since corticosteroids have been described to exert a detrimental effect on tendons, it cannot be determined to which extent chronic systemic inflammation contributes to tendon degeneration [5]. Therefore, the aim of this study was to investigate whether an allergic condition inducing systemic inflammation negatively affects the structure and function of tendons and can promote the development of tendinopathies. To address this question, we employed a mouse allergy model to study the impact on tendon quality and retrospectively analysed epidemiological data from more than 10.000 patients collected during a health study to investigate, if patients suffering from an allergy are at an increased risk to develop a tendinopathy [24].

## Results

Three months old C57/BL6 mice were sensitized with recombinant Timothy-grass pollen allergen Phl p 5 adjuvanted with aluminiumhydroxyde (Alum), followed by a repeated intranasal allergen challenge 11 weeks after the last sensitization (**Fig. 1A)**. For reasons of simplicity, these sensitized and challenged mice will be designated as allergic. To test for potential side effects elicited by the adjuvant, we included a group sensitized with Alum and challenged with PBS. Control animals received PBS both for sensitization and challenge.

**Fig. 1:**
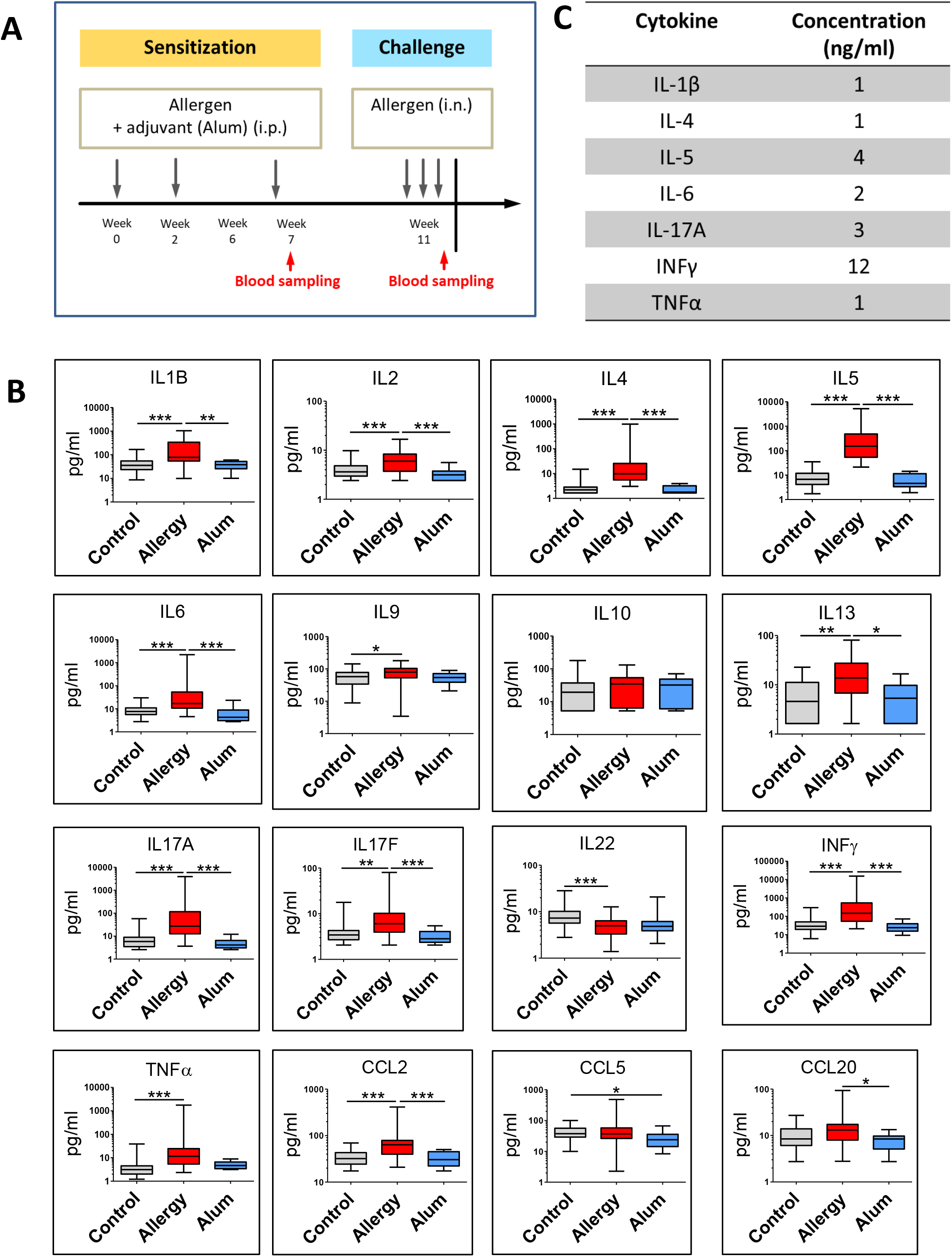
Induction of an allergic response in mice. (**A**) Experimental setup. (**B)** Multiplex bead array analysis reveals significantly elevated serum levels of TH1/pro-inflammatory (IFNγ, TNFα, IL1β, IL2, IL6), TH2/Treg related cytokines (IL4, IL5, IL10, IL13), TH17 cytokines IL17A and IL17F and selected chemokines in the blood of allergic mice compared to control or Alum-treated animals; Data are shown as Mean + SEM, *p<0.05, **p<0.01, ***p<0.001, One-way ANOVA (Kruskal-Wallis and Dunn’s Multiple Comparison Test), n (control/allergy) = 16, n (Alum) = 6. (**C**) Concentrations of selected cytokines used to formulate a cocktail simulating the cytokine concentrations measured in mouse sera.

### Allergic sensitization induces inflammatory serum cytokines/chemokines

To confirm an allergy-induced increase in circulating cytokines/chemokines, a bead based multiplex immunoassay was performed 1 day after the last sensitization step (day 48) and 1 day after the last intranasal challenge (day 77). A clear elevation of the majority of cytokines analysed was already evident after the sensitization phase (**supplement Fig. S1**) and an even more pronounced, significant increase of both Th1 and Th2 cytokines was observed post-challenge, including IL4, IL5, IL13, and IL17, which are considered to be the major contributors to allergy and asthma. No significant difference could be detected between the control and Alum-treated group (**Fig. 1B**). Due to the limited availability of mouse sera needed for the *in vitro* experiments we generated an allergy-like cytokine cocktail by combining cytokines that were at least three times above the control level in “allergic” animals (allergy simulating cytokine cocktail; ASCC). The cytokine concentrations chosen were based on the highest values measured in the blood of allergic animals (**Fig. 1C)**.

### Tendons from allergic animals display altered biomechanical and structural properties

Biomechanical testing revealed that flexor tendons from allergic mice displayed a significantly (p>0.05) reduced tendon stiffness (**Fig. 2C**), whereas no effect could be seen regarding maximum tensile load (**Fig. 2D**). However, taking into account the significantly increased cross sectional tendon area in allergic animals (**Fig. 2B**), both Young’s modulus (**Fig. 2E**) and tensile stress (**Fig. 2F**) were significantly reduced in this group compared to the controls.

**Fig. 2:**
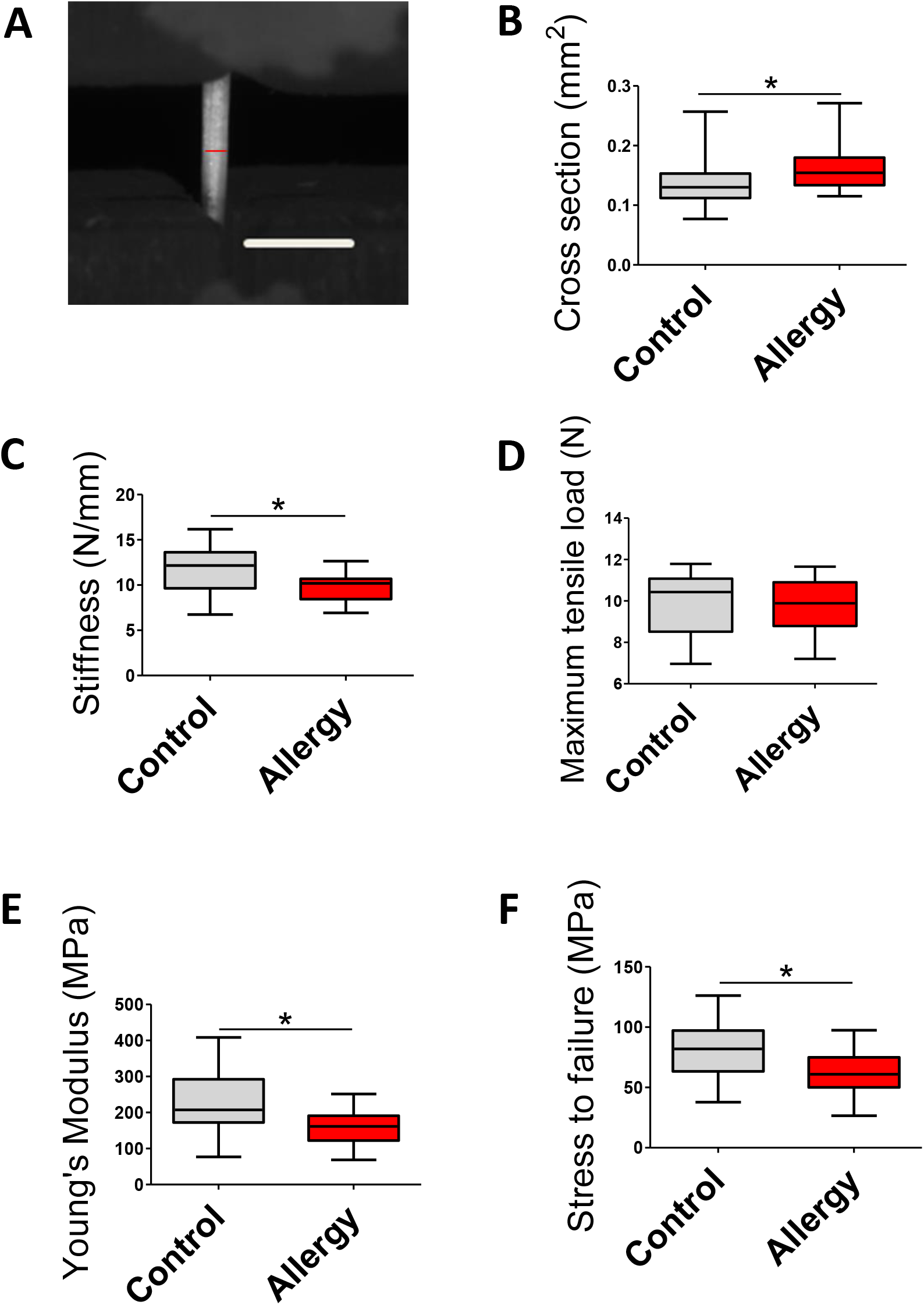
Biomechanical testing parameters. (**A**) Flexor tendon diameters were measured at 0.1N preload using two cameras positioned at an angle of 90° to each other. (**B**) The cross sectional area of tendons from allergic mice was significantly larger than that from control animals; *p<0.05, Mann-Whitney U test, n≥22. (**C)** Tendons from allergic animals displayed a significantly reduced stiffness; *p<0.05, Unpaired 2-tailed *t* test, n≥21). (**D)** No difference in maximal tensile load could be detected; Unpaired 2-tailed *t* test, n≥21. (**E**) Young’s Modulus and (**F**) stress to failure were significantly lower in tendons from allergic mice; *p<0.05, Unpaired 2-tailed *t* test, n≥21. Scale bar: 1mm.

In order to examine whether this observed increase in tendon cross sectional area relates to alterations in the extracellular matrix, we analysed tendons from the control and allergy group both at the light and electron microscopic level (**Fig. 3**). Polarized light microscopy revealed that tendons from allergic animals not only showed a significantly looser collagen packing (**Fig. 3A**), but also a significantly less parallel fibre orientation (**Fig. 3B**).

**Fig. 3:**
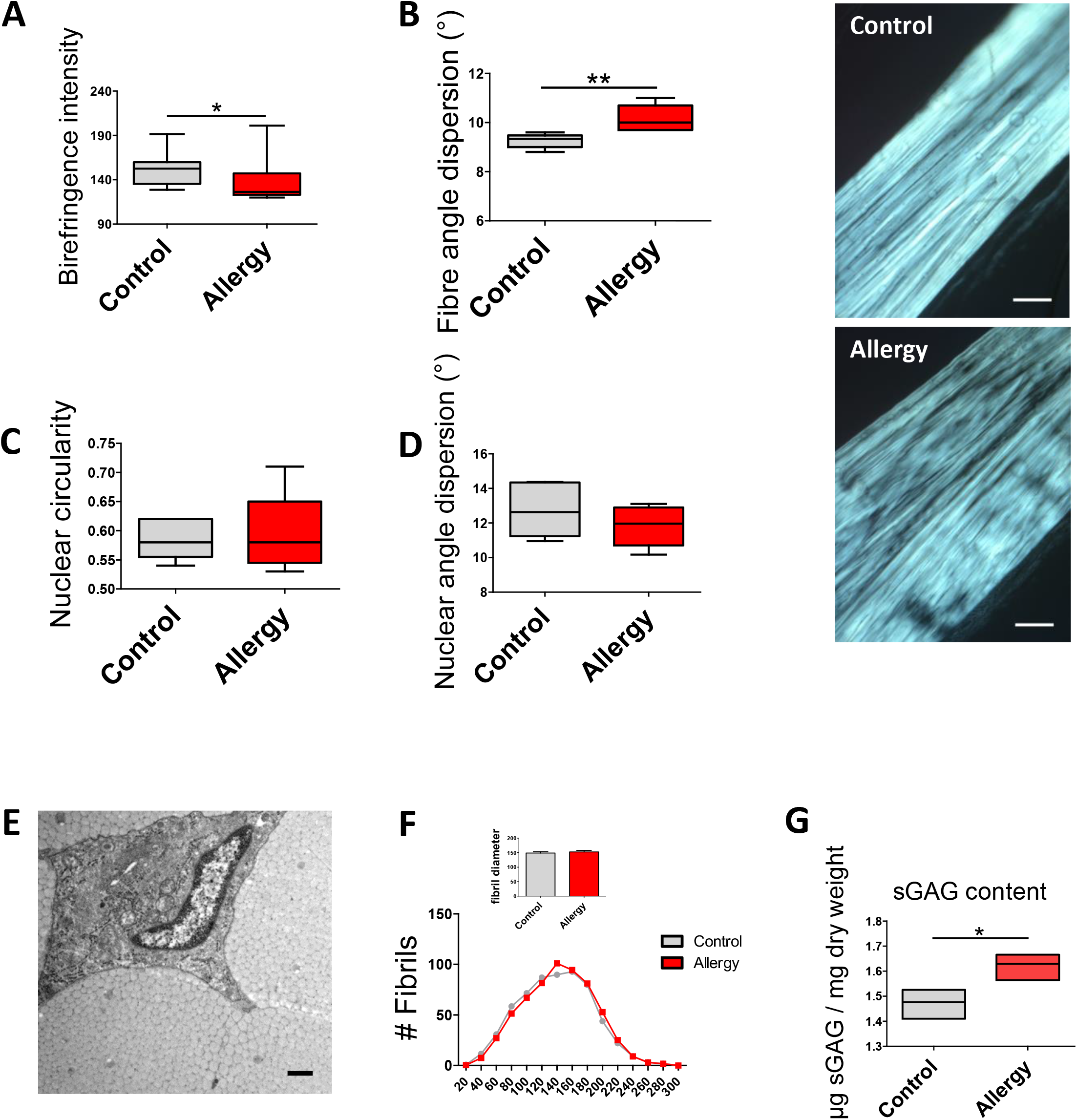
Structural and biochemical analysis of mouse Achilles tendons. (**A**) Examination of histological sections under polarized light revealed a significantly less dense collagen packing in tendons from allergic mice; *p<0.05, Mann-Whitney U test, n≥11. (**B**) Collagen fibers were less parallelly oriented in tendons from the allergy group; *p<0.05, **p<0.01, Mann-Whitney U test, n≥5. (**C, D**) No significant difference between groups could be seen in shape and orientation of cell nuclei; Mann-Whitney U test, n≥5. Scale bar: 100 μm. (**E, F**) Ultrastructural analysis of mouse Achilles tendons. Shown are frequencies of fibril diameters of control and allergic mice determined by TEM (n = 600 fibrils of ≥5 animals each). Mean±SEM are displayed. Analysis of electron microscopic images of Achilles tendons did not show a significant difference in fibril size distribution (insert). The statistical model of Linear mixed model fit by maximum likelihood (‘lmerMod’) was used yielding a p-value of p = 0.55. Scale bar: 500 nm. (**G**) Glycosaminoglycan (GAG) analysis revealed a significantly increased amount of GAGs in tendons from allergic animals; *p<0.05, Unpaired 2-tailed *t* test, n = 3.

In contrast, no differences between the groups could be detected regarding the nuclear aspect ratio and orientation of cell nuclei (**Fig. 3C, D**). The same accounted for fibril diameter distribution as revealed by ultrastructural analysis of the Achilles tendons (**Fig. 3E, F**). Finally, the glycosaminoglycan content of tendons from “allergic” mice was found to be increased compared to tendons from control mice (**Fig. 3G**).

### Allergic sensitization elicits a matrix remodelling phenotype in vitro and in vivo

Tendon-like constructs (**Fig. 4A**) were treated with ASCC for 72 h, changing the media every day to warrant a stable cytokine concentration over the entire incubation time, and the mRNA expression of a variety of matrix-, fibrosis-, and inflammation-related proteins was analysed (**Fig. 4B**). Similarly, we treated tendon-like constructs with pooled sera from “allergic” or control mice for 24h, and performed qPCR analysis targeting the same genes as analysed in the experiments with ASCC (**supplement Fig. S2)**. Of the genes of interest, we specifically focused on those which yielded similar results in both approaches. Interestingly, we found a significantly increased expression of Matrix Metalloproteases 1a (Mmp1a), −2 (Mmp2), −3 (Mmp3), Epiregulin (Ereg) and Interleukin 6 (IL6) in both serum and ASCC-treated 3D-tendon constructs, albeit to a varying extent (**Fig. 4B**). At the protein level we confirmed a significant elevation of MMP1 and MMP2 in tendon-like constructs exposed to ASCC (**Fig. 4C**),.

**Fig. 4:**
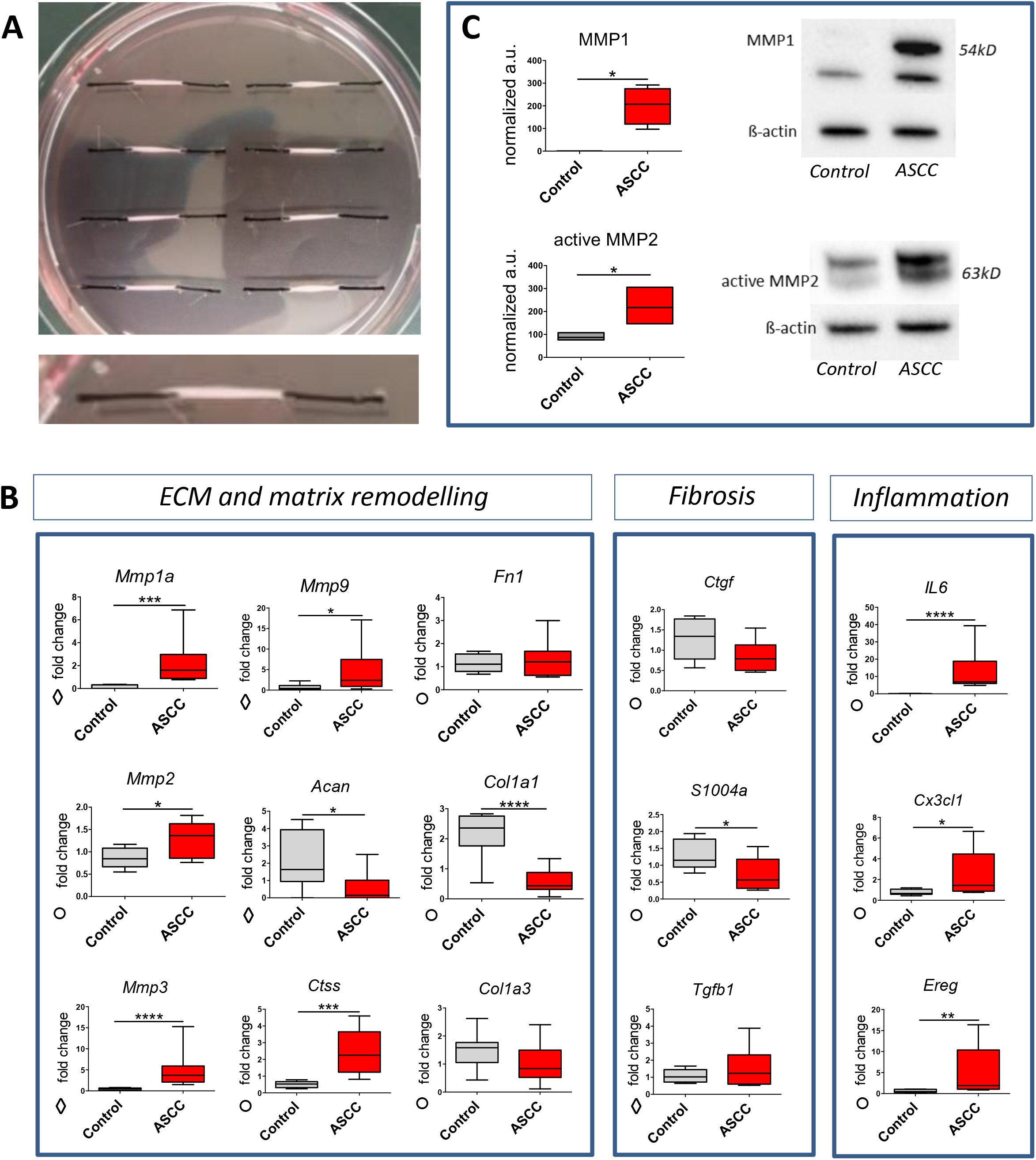
qPCR and Western blot analysis on 3D tendon-like constructs. (**A**) Image of tendon-like constructs spread between silk sutures mounted on a collagen-coated petri dish. (**B**) Treatment of tendon-like constructs with an allergy stimulating cytokine cocktail (ASCC) results in upregulation of selected target genes; *p<0.05, **p<0.01, ***p<0.001, ****p<0.0001, Mann-Whitney U test (◇) or Unpaired 2-tailed *t* test (○) depending on normal distribution, n = 9. (**C**) Western blot analysis of tendon-like constructs exposed to the ASCC shows a significant upregulation of Mmp1 and Mmp2; Bars represent mean ± SEM; *p<0.05, Mann-Whitney U test, n = 4. Scale bar: 1cm.

To assess whether the activity of a matrix metalloprotease is also increased *in vivo*, we analysed the expression of Mmp2 on cryosections of mouse Achilles tendons. Indeed, fluorescent staining revealed a significantly increased expression of Mmp2 in tendons from allergic mice (**Fig. 5A**). Moreover, we found Mmp2 to display both significantly elevated protein levels and enzyme activity as demonstrated by zymography (**Fig. 5B, C**).

**Fig. 5:**
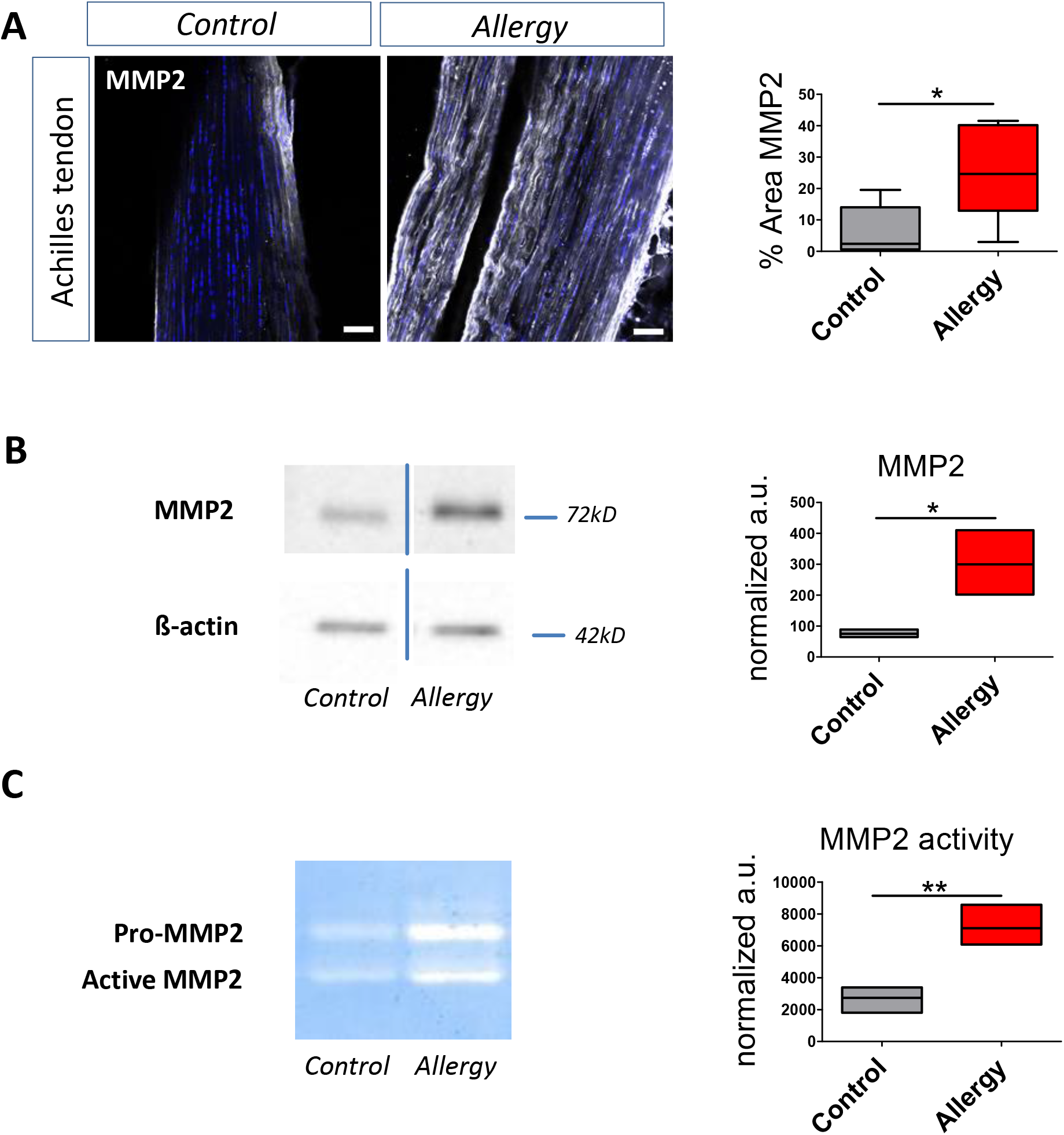
MMP2 expression in mouse Achilles tendons. **(A)** Quantification of immunofluorescence staining on cryosections reveals a significantly increased MMP2 expression in tendons from allergic mice; Representative image and quantification; *p<0.05, Mann-Whitney U test, n ≥ 5. (**B**) Western blot analysis shows upregulation of MMP2; Representative image and densitometric quantification, normalization to β-actin; *p<0.05, Unpaired 2-tailed *t* test, n = 3. (**C**) MMP2 activity is upregulated in allergic animals as shown by gelatin zymography; representative image and densitometric quantification, normalization to ß-actin; **p<0.01, Unpaired 2-tailed *t* test, n = 3. Scale bar: 50 μm.

### Allergic animals show increased macrophage activation in tendons

As we had observed an increase in mRNA levels encoding Ereg in ASCC-treated 3D tendon-like constructs and we previously had identified a cell population (“tenophages”) expressing Ereg and macrophage-associated marker proteins [25], we next examined if increased levels of circulating cytokines affects immune cell marker expression within the tendon tissue. Indeed, increased expression of F4/80, a marker highly and constitutively expressed on most resident tissue macrophages, of CD68/macrosialin, which is a heavily glycosylated glycoprotein highly expressed in macrophages and other mononuclear phagocytes, of Ionized calcium binding adaptor molecule 1 (Iba1), a microglia/macrophage-specific calcium-binding protein, and of CD163, a specific marker for monocytes/macrophages (cell populations exhibiting strong inflammatory properties), was evident in cryosections prepared from Achilles tendons of allergic mice (**Fig. 6**).

**Fig. 6:**
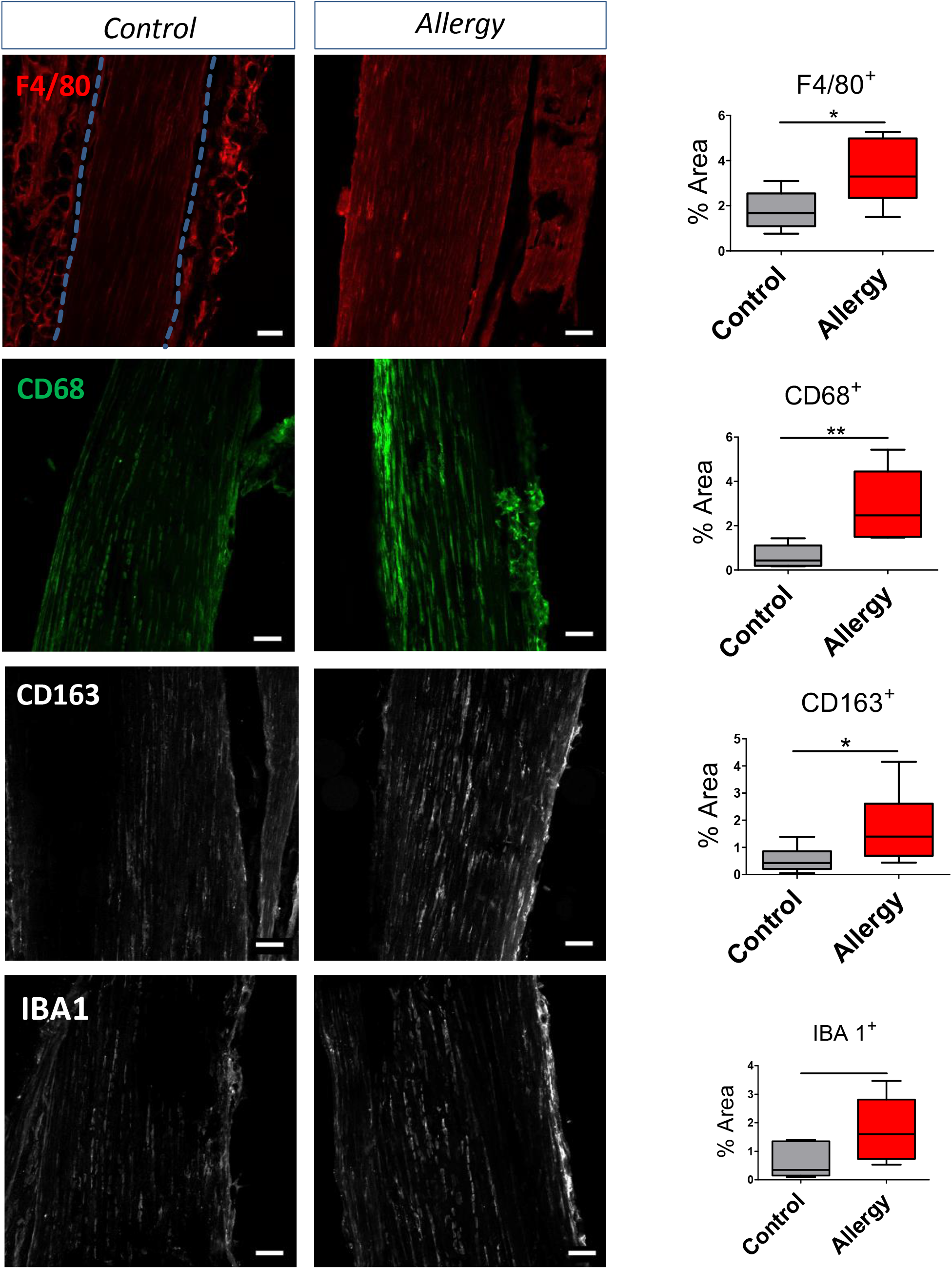
Macrophage marker expression *in vivo*: Achilles tendons from allergic mice show a clearly increased expression of (**A**) F4/80, (**B**) CD68, (**C**) CD163, and (**D**) Iba-1 (p=0.0519) compared to control animals as evidenced by fluorescent staining of cryosections. Representative images and densitometric quantification are shown; *p<0.05, Mann-Whitney U test, n ≥ 6. Scale bar: 50 μm.

Given this increase in expression of immune cell-related markers within allergic tendons, we speculated whether this might correspond with an increased activation state. To address this question, we stained cryosections from control and allergic animals and 3D-tendon like constructs which had been incubated with or without the ASCC for markers indicative of macrophage activation such as the lipid droplet associated protein perilipin. Interestingly, we found a significantly elevated expression of peripilin 1 both *in vivo* and *in vitro* (**Fig. 7A, B**).

**Fig. 7:**
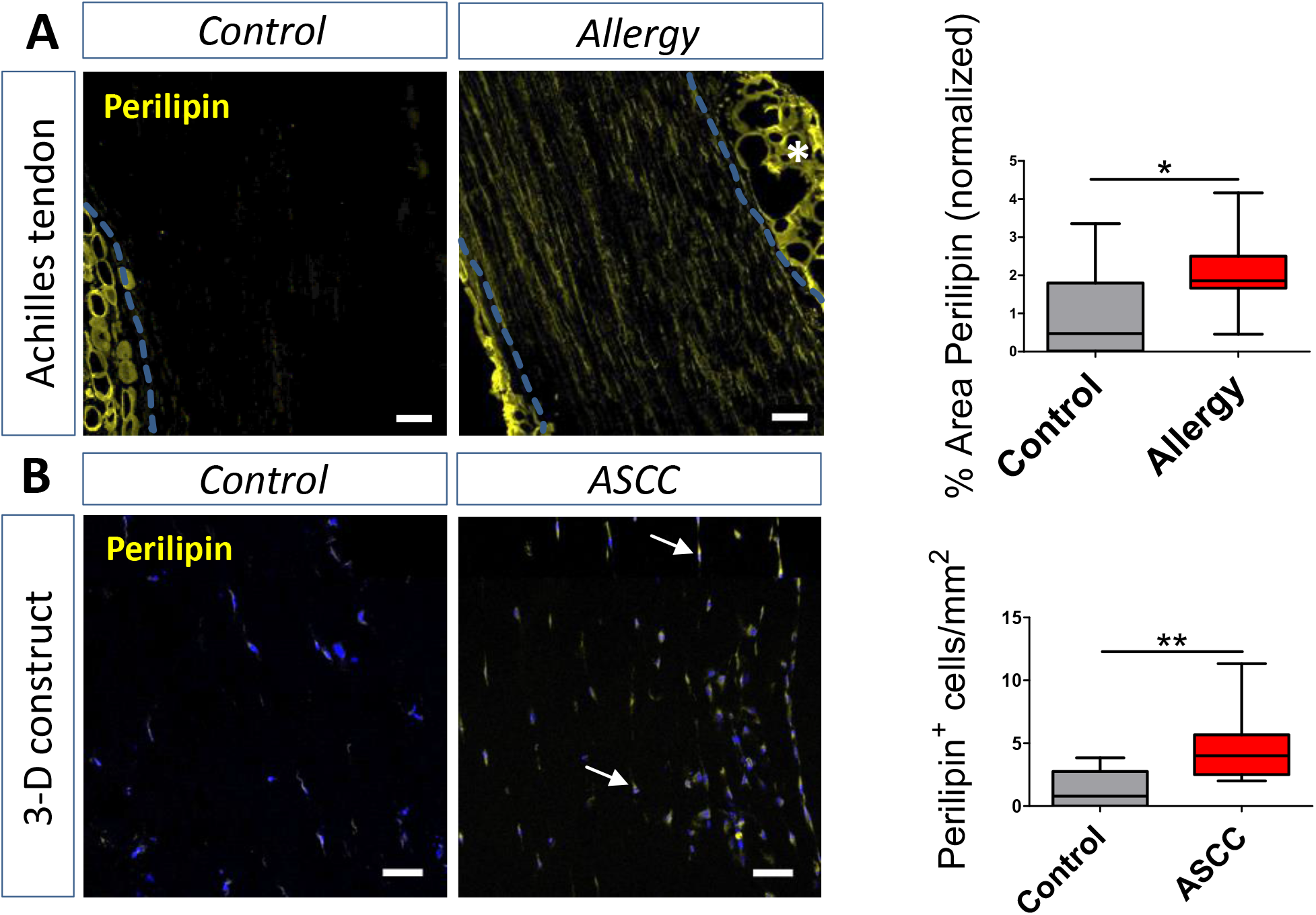
Perilipin expression *in vivo* and *in vitro*. **(A)** Perilipin is significantly higher expressed in Achilles tendons from allergic mice; representative image and quantification of perilipin expression, normalized to control settings; *p<0.05, Unpaired 2-tailed *t* test, n ≥ 11. (**B**) *In vitro* the number of perilipin positive cells in 3D tendon-like constructs exposed to ASCC is increased compared to controls; **p<0.01, Mann-Whitney U test, n ≥ 8. Blue DAPI stain indicates cell nuclei. White arrows point toward perilipin positive cells. White asterisk indicates adipose tissue. The dotted blue line demarcates the tendon proper. Scale bar: 100 μm.

### Allergy is a moderate risk factor for tendinopathy

Based on the data from a large health survey study conducted in the state of Salzburg (Paracelsus 10.000 study), retrospective statistical analysis of patients’ records regarding a potential relationship between allergic conditions and reported tendon-related diseases was performed. Overall our sample consisted of observations from 10043 study participants aged between 40 and 75 years. For comparability between the different logistic regression models applied (see Methods section), we used only patient data sets which included information for all variables included in multivariable logistic regression model 3. This dataset consisted of data from from 8471 patients. To test for the validity and plausibility of our approach, we further analysed patients’ data focussing on risk factors known to be associated with tendinopathies (e.g overhead work/impingement, rheumatoid arthritis, diabetes, obesity, smoking). Baseline characteristics of study participants and the proportions of persons displaying the above mentioned risk factors are listed in Table 1.

**Table 1:**
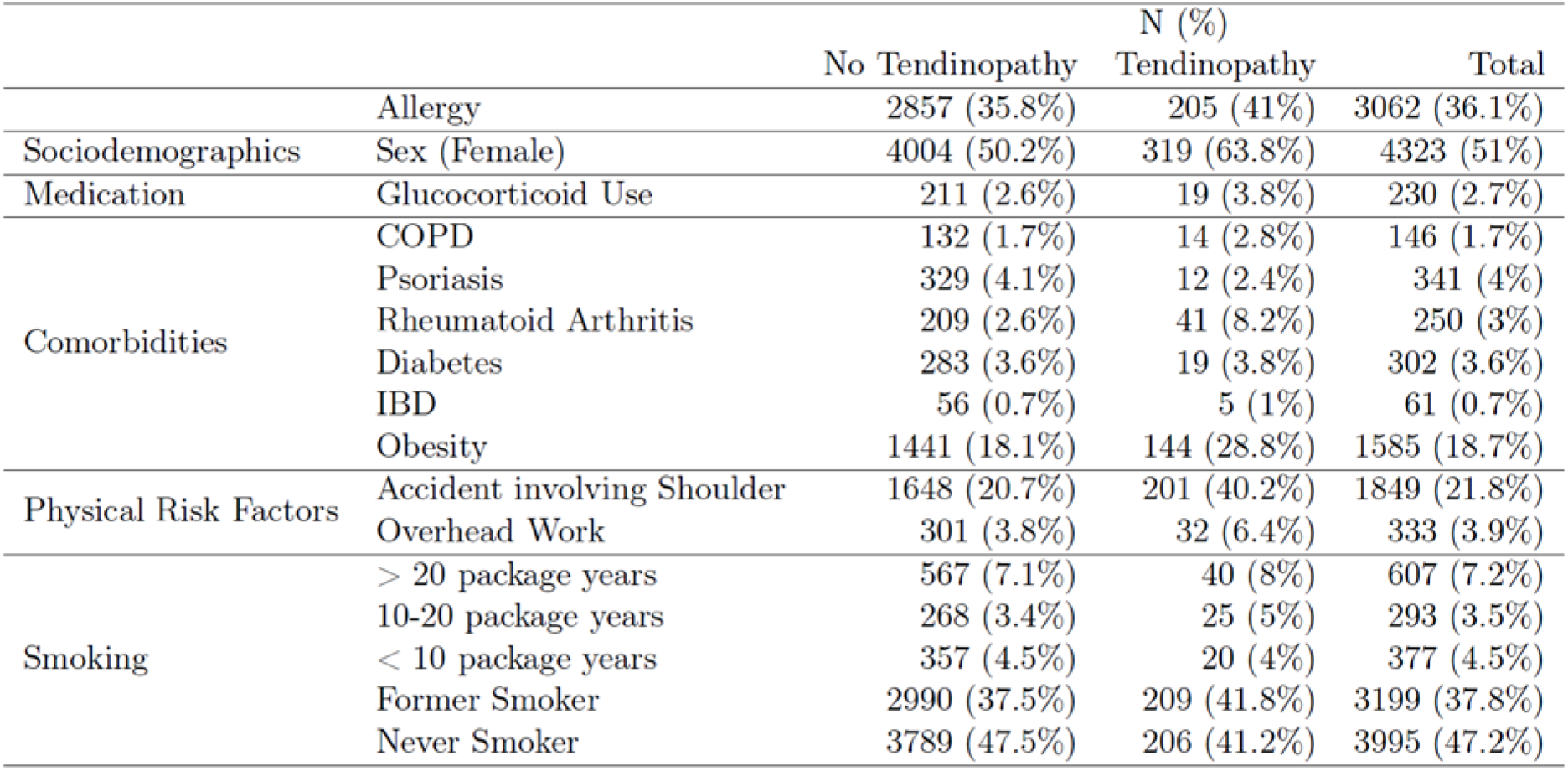
Baseline characteristics of study participants.

Surprisingly, more than a third of all participants were affected by some form of allergic symptoms (**Table 1**). About 20% of all persons had an accident involving the shoulder and about 4% performed overhead work. The proportion of obese persons in the study population was 18.7%, whereas the number of persons affected by other tendinopathy-associated risk factors ranged between 0.7% and 4%.

To investigate if there is a correlation between allergies and signs of tendinopathy, the outcome variable was defined as follows: participants of the health survey needed to state at least one of the following terms: nocturnal shoulder pain OR inability to lift a milk carton overhead OR declaration of a tendinopathy-related keyword, which applied to 2355 participants (**Table 2**). A more restricted outcome variable defining tendinopathy as the presence of a tendinopathy-related keyword OR a combination of nocturnal shoulder pain AND the inability to lift a milk carton was also employed, however this criterion only applied to 500 patients and is reported in the supplementary information (**Table S2**).

**Table 2:**
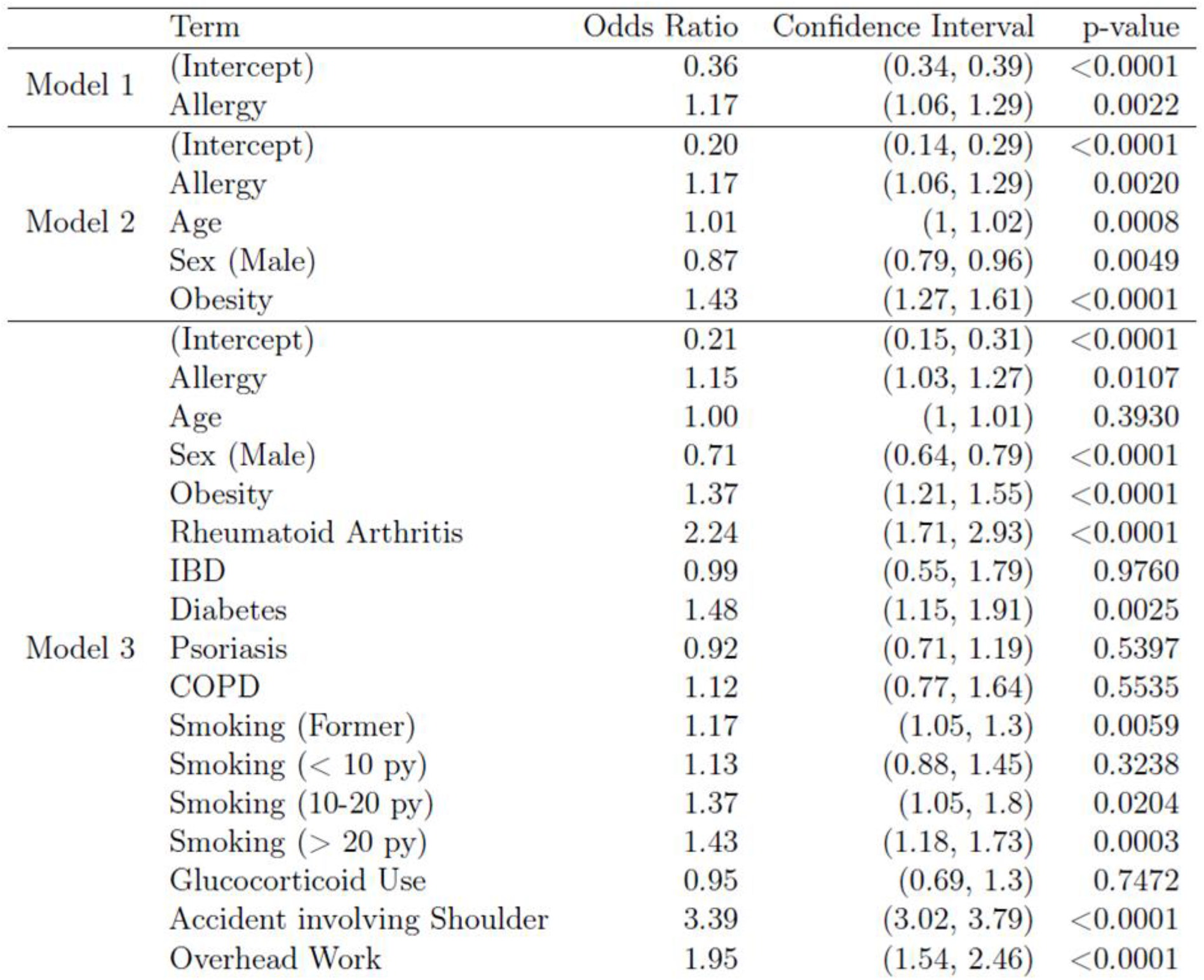
Odds Ratios, 95% Confidence Intervals for the Odds Ratio and p-values for all terms in the three models when the outcome is presence of a tendon surgery OR inability to lift a carton of milk overhead OR nocturnal shoulder pain.

Table 2 and Table S2 show the odds ratios (OR) for the predictor variable “allergy” in three different logistic regression models, the odds ratio being a statistical measure indicating the strength of a relationship between two characteristics. The logistic regression analysis yielded an OR for the allergy term lying between 1.17 and 1.15, respectively, depending on how confounding effects have been controlled for. When correcting for confounders known to increase the risk of developing a tendinopathy, an OR of 1.15 was determined (*P=0.0107*), indicating that the odds to develop a tendinopathy is 1.15 times higher compared to non-allergic persons. Among the confounder variables known to be risk factors for tendinopathies rheumatoid arthritis (OR=2.24), overhead work (OR=1.95), diabetes (OR=1.48), obesity (OR=1.37), and smoking (OR ranging between 1.17 and 1.43 depending on pack years) also revealed elevated odds ratios. Interestingly, moderate smoking seems to have a similar effect as the presence of an allergy.

## Discussion

Tendinopathies still present a major medical challenge for physicians and orthopaedics as treatment options are limited, not at least due to an incomplete understanding of their aetiology. Major risk factors, apart from repetitive overload, known so far include diabetes, obesity, several autoimmune diseases and smoking [26]. Since there is evidence that persons suffering from diseases associated with a systemic low-grade inflammation run a higher risk of developing tendinopathies or sustaining a tendon rupture, we were interested to elucidate whether the presence of an allergy would be sufficient to alter the structure and function of tendons [27]. In the present study, we therefore investigated the impact of a systemic inflammation induced by timothy grass pollen allergy in a mouse model [24], the advantage of this animal model being that no additional, potentially confounding factors (e.g. age, obesity, other autoimmune diseases) mask the effects induced by the allergic inflammation. Respiratory exposure to the grass pollen allergen resulted in an increase in systemic levels of key cytokines known to be associated with allergic diseases - IL4, IL5, IL13, and IL17 [28–30]. Three of them (IL4, IL5, IL17A) were included into an allergy simulating cytokine cocktail (ASCC) together with TNFα, IL1β, INFγ, and IL6. All the cytokines included were either least three-fold higher in the blood sera of allergic animals compared to control animals, or have been shown to be significantly elevated in the blood of allergic children [31].

On the functional level, biomechanical testing revealed a significant decrease in both Young’s modulus and tensile stress in the allergy group, the cross sectional area being significantly increased compared to the control group. Interestingly, similar findings were seen in a study by Hernandez et al. in which they showed that severe burn-induced systemic inflammation led to a decrease in stiffness and ultimate force of rat Achilles tendons [32]. Moreover, the authors report a decrease in organized collagen fibers 14 days after a burn, which is in agreement with our observation of looser collagen packing and less parallel oriented fibers in tendons from allergic mice. As shown by ultrastructural analysis, the increase of tendon cross sectional area in the allergic animals is not a consequence of larger collagen fibril diameters. However, glycosaminoglycan (GAG) content in the flexor tendons of allergic animals was significantly increased. This elevated level of sulfated GAGs might lead to increased tissue swelling accounting for the change in size, as well as be related to the observed decrease in elastic modulus and tensile stress, since there is evidence for a causal GAG-biomechanics relationship in diseased tendons. In an equine tendon injury model it has been shown that injured tendons displayed both a significantly decreased modulus and ultimate tensile strength with a concomitant increase in glycosaminoglycan content throughout the tendon [33], which is very well in line with our findings. Moreover, a significant increase in sulfated glycosaminoglycan content in human patellar tendinopathy has been described [34, 35].

In order to shed light on potential underlying mechanisms accounting for the observed structural changes, we investigated the effects of both mouse sera and cytokine cocktail on the expression of a variety of genes in 3D-collagen embedded tendon cells *in vitro*, focusing predominately on matrix remodelling and inflammation-related proteins. With both approaches we found a significant mRNA upregulation of matrix remodelling enzymes Mmp1, Mmp2 and Mmp3, as well as of IL6, a key cytokine known to be involved in the pathogenesis of several chronic inflammatory diseases, and of epiregulin, shown to be increased upon inflammatory stimulation in macrophage-like tendon cells [36]. The difference in expression levels in response to the treatment can likely be explained by differences in incubation time (24 vs. 72 hours), the shorter incubation time owing to a limited amount of serum, and differences in cytokine composition, the ASCC containing only a selected number of cytokines, whereas the serum applied not only comprises the cytokines measured by multiplex bead array, but furthermore a plethora of components (including chemokines, cytokines, and lipid mediators) potentially enhancing or counteracting the effect of the proteins analysed.

The observed up-regulation of MMP2 on the protein level both of lysates of tendon-like constructs exposed to ASCC and in Achilles tendons from allergic mice potentially contribute to the structural changes seen in these tendons, given that an increased expression of MMP1 and MMP2 has also been reported in human tendinosis tissue [37–39].

As reported in our previous study, mouse tendons harbour a sub-population of tendon cells expressing immune cell-related markers and appear to possess phagocytic activity similar to macrophages [36]. Here, we show that these markers including F4/80, CD68, and CD163 are significantly increased in Achilles tendons from allergic mice compared to control animals. Since a high expression of CD163 in macrophages is characteristic for tissues responding to inflammation and enhanced presence of CD163+ macrophages with an increase in soluble CD163 (sCD163) has been described in a variety of human inflammatory diseases, including asthma, rheumatoid arthritis and diabetes, the increased expression of CD163 observed in allergic animals is consistent [40, 41]. Also, the higher expression level of IBA 1 is in line with previously published results, demonstrating that IBA1 is significantly upregulated in mice upon stimulation with IL1ß/TNFα [42]. The altered macrophage/ tenophage marker profile is indeed also reflective of alterations in their activation state, since we found that tendons from allergic animals as well as ASCC-treated 3D tendon constructs *in vitro* displayed a significantly elevated expression of perilipin 1. This protein is not only present on the surface of adipocyte lipid droplets (LDs), but has also been reported to be expressed by activated macrophages, as lipid accumulation in phagocytes is a hallmark of infection and sterile inflammation [43, 44]. These changes in immune cell function, such as myeloid cell activation, have been described to be connected to profound changes in LD numbers and LD protein composition. Thus, these lipid-rich, cellular organelles appear to have a vital role in antigen cross-presentation, interferon (IFN) responses, production of inflammatory mediators, and pathogen clearance [45].

Overall, the present study demonstrates that the presence of an allergy-induced systemic inflammation over a relatively short period of time is sufficient to have a detrimental effect on tendon quality and function. The relevance of these findings *in vitro* and *in vivo* was further underpinned by the analysis of data obtained from a large health survey (Paracelsus 10.000 study). The association between low level systemic chronic inflammation and the occurrence of tendinopathies has been reported in several studies [9, 46]. Importantly, systemic inflammation accompanies also many conditions known to be associated with the development of tendinopathies, such as performing heavy overhead work, suffering from diabetes or smoking. To benchmark our search strategy to define the outcome variable “tendinopathy” (presence of a surgery OR inability to lift a carton of milk OR nocturnal shoulder pain), we probed the available cohort for the above mentioned risk factors (Rheumatoid arthritis, obesity, diabetes, smoking, overhead work). All of them presented with a significantly elevated odds ratio, which are well within the range of odds ratios reported by other studies [26, 47–49].

The Paracelsus 10.000 study is a prospective cohort study and did not specifically and exclusively focus on tendons and ligaments. This means that tendinopathy has not explicitly been clinically diagnosed by a physician based on ultrasound analysis, but tendon/ligament-related surgeries and tendon-related pain conditions have been self-reported by participants. The data analysed for the present study are of retrospective and cross sectional nature. Due to the difficulty to discriminate between traumatic and acute on chronic injuries, we decided to focus on tendinopathy-related symptoms including nocturnal shoulder pain, the inability to raise the arm above 90° and patient-reported keywords such as calcification, tenosynovitis and carpal tunnel syndrome and deliberately excluded all reports on most likely acute tendon or ligament tears, as well as terms which could not be assigned unequivocally. Model 3 represents the most stringent way to analyse the data set controlling for a potential effect of glucocorticoid use on tendons while avoiding collider bias by also controlling for other comorbidities which are known to affect tendons and for which glucocorticoids could be prescribed yielding an OR for allergy of 1.15 (p=0.0107). In humans, allergy is so far an unrecognized risk factor for tendon disease, an insight which might have implication for future prevention and/ or treatment strategies.

### Summary

In the present study conducted in a mouse model, we demonstrate that an allergy-induced systemic inflammation impacts on tendons, both on the functional as well as on the structural level. Analysis of patients’ records from a health survey study suggest that the presence of an allergic condition accompanied by a systemic inflammation also affects tendons in human and might be considered a risk factor for the development of tendinopathies (**Fig. 8**).

**Fig. 8:**
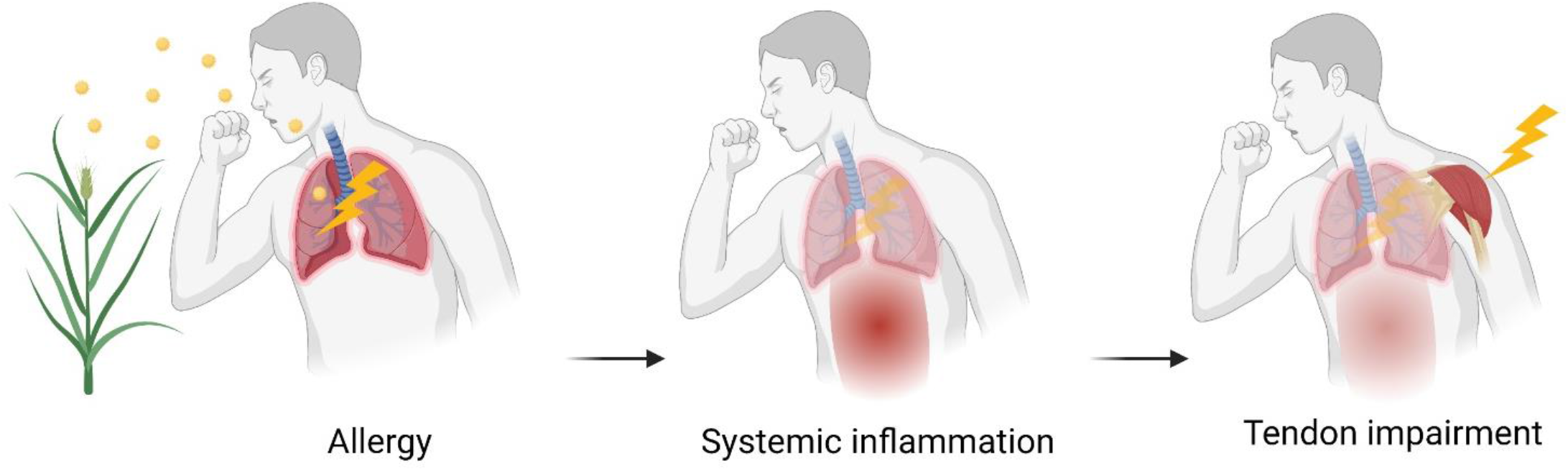
Graphical summary

## Materials & Methods

### Animals

Female C57BL/6 mice (aged 10–12 weeks, 20-25g) were purchased from Charles River Laboratories, Germany. All animals were acclimatized to standard laboratory conditions (14-h light, 10-h dark cycle) and given free access to rodent chow and water at the animal facility at the Paracelsus Medical University, Austria. All experimental procedures were approved by the Austrian Ministry of Science (BMWFW-66.019/0023-WF/V/3b/2017) and carried out in compliance with national ethical guidelines.

### Allergy Induction

Mice were rendered allergic by exposing them to a plant allergen according to a recently published immunization regimen [24]. Briefly, recombinant Phl p 5.0101 (Phl p 5) was purchased from Biomay AG. The animals were divided into three groups: controls (*n* = 55), allergy model (*n* = 55), and adjuvant group which received aluminiumhydroxide (Al(OH)3) only (*n*=6). The control group received all treatments using only the vehicle solution (phosphate-buffered saline, PBS). Animals of the allergy group were immunized intraperitoneally (i.p.) with 1 μg Phl p 5 adjuvanted with Al(OH)_3_ (Alu-Gel-S from Serva) in PBS (50% v/v, total volume: 200 μl) at weeks 1, 2, and 7. In week 11, starting 4 days before the perfusion (day 75), this group was challenged three times with a daily dose of 5 μg Phl p 5 in 40 μl PBS intranasally (i.n.; on days 71, 74 and 75). During this procedure, all mice (including the control and adjuvant group) were briefly anesthetized with isoflurane.

### Analysis of Blood Parameters

Blood samples were taken twice, once after the last sensitization step (day 48) and another time at the ending of the experiment (day 76), and incubated for 1 h at room temperature. After centrifugation (10 min), the sera were collected and stored at −80°C until measurements. Serum levels of a selective panel of cytokines and chemokines (IL1β, IL2, IL4, IL5, IL6, IL9, IL10, IL13, IL17A, IL17F, IL22 INFγ, TNFα, CCL2, CCL5, CCL20) were measured with BioLegend’s LEGENDplex™ bead-based immunoassays (Biolegend, San Diego, U.S.) according to the manufacturer’s instructions.

In order to mimic allergic conditions, we used a cocktail containing a number of cytokines which have turned out to be most significantly elevated upon allergy-induction and at least three-fold elevated above control values (Fig. 1B). This allergy-simulating cocktail used for *in vitro* experiments comprised 7 different cytokines (IL1β, IL4, IL5, IL6, IL17A, INFγ, TNFα).

### 3D Tendon-like constructs

In order to better mimic the tendon’s natural environment, we performed most of our experiments using 3D-collagen embedded tendon cell cultures. These artificial tendon-like constructs were established as described previously [50]. In brief, 2.5 x 10^5^ mouse Achilles tendon-derived cells (passage 2) isolated of at least 6 animals per preparation were mixed with collagen type I (PureCol™ EZ Gel solution, # 5074, Sigma-Aldrich, Vienna, Austria; endconcentration 2mg/ml) and spread between two silk sutures pinned with insect pins in rows on SYLGARD 184 (Sigma-Aldrich) coated petri dishes. To improve formation of the constructs, Aprotinin, Ascorbic acid, and L-Proline were added to the cell culture medium. After contraction of constructs over the course of 12 days, a cytokine cocktail containing the respective cytokine concentration measured in the sera of allergic mice (1ng/ml IL1β, 1ng/ml IL4, 4ng/ml IL5, 2ng/ml IL6, 3ng/ml IL17A, 12ng/ml INFγ, and 1ng/ml TNFα) was added to the culture medium. Medium changes and addition of fresh cytokines was performed every day. After incubation for 72 hours, constructs were harvested and stored either in TRIReagent (Sigma-Aldrich, Austria) for further qPCR analysis, fixed in 4% paraformaldehyde for immunohistochemical analysis or frozen at −80°C for subsequent western blot analysis. TNFα, INFγ, and all interleukins were purchased from PeproTech (Vienna, Austria) (supplement Table S3).

### Preparation of tissue sections

Mouse Achilles tendons and mouse 3D tendon-like constructs were fixed in 4% paraformaldehyde for 12 hours at 4 °C, and after several washes in phosphate-buffered saline (PBS) and cryo-preservation in 30% sucrose in PBS were embedded in cryomedium (Surgipath Cryogel^®^, Leica Microsystems, Vienna, Austria). Subsequently, 12 μm cryosections were prepared using a Leica CM1950 cryostat.

### Immunohistochemistry

Immunohistochemical detection of immune cell-related markers was performed on cryosections of tendons and 3D tendon-like constructs, respectively. After a 5 min rinse in Tris-buffered saline (TBS; Roth, Karlsruhe, Germany) slides were incubated for 1h at room temperature (RT) in TBS containing 10% donkey serum (Sigma-Aldrich, Vienna, Austria), 1% bovine serum albumin (BSA; Sigma-Aldrich, Vienna, Austria), and 0.5% Triton X-100 (Merck, Darmstadt, Germany). Followed by a 5 min rinse, slides were subsequently incubated for double or triple immunohistochemistry (overnight at 4°C) with antibodies directed against EGF-like module-containing mucin-like hormone receptor-like 1 (F4/80, MCA497RT, Serotec, Oxford, UK; 1:100), Cluster of Differentiation 68 (CD68, #sc20060, Santa Cruz, Dallas, USA; 1:50), ionized calcium-binding adapter molecule 1 (IBA1, #019-19741, FUJIFILM Wako Pure Chemical Corporation; 1:100), Cluster of Differentiation 163 (CD163, #ab182422, Abcam, Cambridge, UK; 1:100), and MMP1 (Proteintech, Rosemont, USA, #10371-2-AP), all diluted in TBS, BSA, and Triton X-100.

After a rinse in TBS (four times 5 min) binding sites of primary antibodies were visualized by corresponding Alexa488-, Alexa568-, or Alexa647-tagged antisera (1:500; Invitrogen, Karlsruhe, Germany) in TBS, containing 1% BSA and 0.5% Triton X-100 (1h at RT) followed by another rinse in TBS (four times 5 min). Some of the slides received an additional nuclear staining using 4’,6-Diamidino-2-phenylindol dihydrochloride (DAPI). For that, slides were incubated 10 min (1:4000, stock 1 mg/ml, VWR, Vienna, Austria) followed by rinsing in PBS (three times 5 min). All slides were embedded in Fluoromount^™^ Aqueous Mounting Medium (Sigma Aldrich, Vienna, Austria). Negative controls were performed by omission of the primary antibodies during incubation and resulted in absence of immunoreactivity.

### Confocal imaging

Confocal imaging was performed using a LSM1 700 confocal microscope (Zeiss) equipped with 405 nm (5 mW fiber output), 488 nm (10 mW fiber output), 555 nm (10 mW fiber output) and 639 nm (5 mW fiber output) diode lasers, a main dichroic beam splitter URGB and a gradient secondary beam splitter forLSM 700 using a 10x EC Plan-Neofluar (10x/0.3) or a 20x Plan-Apochromat (20x/0.8) objective (Zeiss, Munich, Germany). Image acquisition was done with ZEN 2010 (Zeiss), and image dimensions were 1024×1024 pixels with an image depth of 16 bit. Two times averaging was applied during image acquisition. Laser power and gain were adjusted to avoid saturation of single pixels. All images were taken using identical microscope settings based on the secondary antibody control stainings.

### Biomechanical testing

For biomechanical testing, digital flexor tendons were isolated from control (n = 55), and allergy-induced (n=55) mice. Immediately after isolation, tendons were wrapped in gauze soaked with PBS to prevent drying and stored at −20 °C until testing. Specimens were tested on a universal material testing machine (Zwick/Roell 500N Zwicki, Ulm/Einsingen, Germany). After application of a preload of 0.1N samples were loaded at a speed of 0.1 mm/min until failure. Stiffness was calculated from the linear proportion of the force/elongation curve and is expressed as N/mm. Tendon material properties were calculated by converting the load-displacement data into stress-strain data using the tendon diameter measured and calculated by using two cameras located at an angle of 90° to each other. Ultimate stress was defined as the stress at tendon failure. Elastic modulus was calculated as the slope of the linear region of the stress-strain curve. Samples, which had partially slipped during tension testing were excluded from the analysis.

### Protein lysates, SDS-PAGE and Western Blot

Tendon-like constructs were washed with ice cold PBS and lysed in RIPA buffer (Sigma-Aldrich, #R0278) supplemented with both a protease cocktail Sigma-Aldrich, #P8340) as well as phosphatase inhibitor cocktail 3 (Sigma-Aldrich, #P0044). Five individual protein lysate preparations were made.

Achilles tendons dissected from at least three individual animals for each group were flash frozen in liquid nitrogen. Following homogenization with a mortar and pestle, ground tissue was resuspended in lysis buffer, incubated on a shaker for 15 min on ice and centrifuged at 13.000 g for 15 min at 4°C to remove cell debris. At least three individual protein lysates were prepared for each group.

Ten to 15 μg of total protein were separated on 10–12% SDS-polyacrylamide gels in Laemmli buffer. Proteins were then transferred to a PVDF membrane (Biorad, Munich, Germany) using 15.6 mM Tris base, 120 mM glycine, and 20% methanol for 1.5 h at 90 V and 4°C. Membranes were blocked in 5% non-fat dry milk powder or 5% BSA hydrolysate in TBS with 0.5% Tween-20, respectively over night at 4°C. Immunodetection was performed using primary antibodies recognizing Mmp1 (Proteintech, #10371-2-AP), Mmp2 (Proteintech #66366-1-lg), Mmp3 (Proteintech #66338-1-Ig), ß-Actin (Sigma #A2228), and secondary horseradish peroxidase (HRP)-labelled goat anti-rabbit antibodies, respectively (BioRad, Munich, Germany). Bands were visualized using the Clarity^™^ Western ECL substrate from BioRad (#170-5060). Band intensities of at least 3 individual experiments were measured densitometrically and normalized to whole protein using the Image Lab Software 5.1 from BioRad (Biorad, Munich, Germany).

### GAG analysis

To obtain a quantifable amount of tendon material, we had to pool flexor tendons from at least 10 animals per group. The content of sulfated glycosaminoglycan of tendons from control and „allergic” mice was assessed by using the Biglycan GAG assay from Biocolor (Carrickfergus, UK) following the manufacturer’s instructions using a spectrophotometer (TECAN Spark, Männedorf, Switzerland). The measurement was repeated three times and the total GAG amount measured was normalized to tendon dry weight.

### Transmission electron microscopy

For ultrastructural analysis, Achilles tendons from control, and „allergic” mice (6 animals per group), were dissected out and processed for transmission electron microscopy as previously described with minor modifications. In brief, samples were incubated in primary fixative (100 mM sodium phosphate buffer pH 7.0 containing 2% glutaraldehyde for 30 min at RT, then transferred to fresh primary fixative and incubated for another 2 h at 4 °C. After several washes in 200 mM sodium phosphate buffer [pH 7.0], specimens were post-fixed in secondary fixative (50 mM phosphate buffer pH 6.2 containing 1% glutaraldehyde and 1% osmium tetroxide; Electron Microscopy Sciences, Hatfield, USA) for 40 min at 4°C. Subsequently, specimens were thoroughly washed with distilled water and en bloc stained in 1% aqueous uranyl acetate for 16 h at 4°C. Samples were dehydrated in graded ethanol series (30%, 50%, 70%, 80%, 95%) followed by four changes of 100% ethanol for 15 min each at RT. Samples were then gradually infiltrated with a mixture of low viscosity resin (UltraBed Kit, Electron Microscopy Sciences) and ethanol according to the manufacturer’s instructions. Finally, samples were placed in embedding moulds filled with pure resin and polymerized at 60 °C for 24 h. Ultrathin sections (80 nm) were cut on a Reichert-Jung Ultracut ultramicrotome (Leica Microsystems, Vienna, Austria). Sections were mounted on Formvar coated 75-mesh copper grids, contrasted with aqueous solutions of uranyl acetate and lead citrate, post-stained with 0.5% uranyl acetate and 3% lead citrate and analyzed with a Zeiss EM 910 electron microscope equipped with a Troendle sharp:eye 2 k CCD camera (Carl Zeiss GmbH, Oberkochen, Germany.

To assess differences in the collagen fibril size distribution between the groups we measured the diameter of 600 fibrils from each animal and classified them into 16 size categories starting with 20-39 nm and progressing in 20 nm steps to the last 320-339 nm (using at least 3 random images per sample).

### Quantitative RT-PCR

Total RNA was isolated from tendon-like constructs (n=4 animals, 3 constructs each) using TRI^®^ Reagent (Sigma-Aldrich; Vienna, Austria) according to the manufacturer’s protocol. RNA yield was quantified using a Nanodrop 2000C (ThermoFisher Scientific, Vienna, Austria) and RNA integrity was verified using an Experion Automated Electrophoresis system (Biorad, Munich, Germany). A minimum requirement of the RNA quality indicator (RQI) >7.5 was chosen.

qRT-PCR was performed as described by Lehner et al. using TaqMan^®^ assays from IDT (Integrated DNA Technologies, Coralville, IA, USA) targeting all genes listed in Table 3 [51]. Amplification conditions were 50 °C for 2 min, 95 °C for 10 min, followed by 40 cycles of 95 °C for 15 s and 60 °C for 1 min. All samples were run in duplicate. CQ values were analyzed using qBasePlus v. 2.4 (Biogazelle NV, Zwijnaarde, Belgium) and normalized relative quantities were calculated by normalizing the data to the expression of previously validated endogenous control genes as described by Vandesompele et al. [52]. As housekeeping genes eukaryotic translation initiation factor 2B subunit alpha (*Eif2b1*), pumilio RNA-binding family member 1 (*Pum1*), and TATA box binding protein (*Tbp*) were used. The normalized quantities were then determined for the candidate genes scaled against the expression values determined for the controls to generate fold changes in expression.

**Table 3.**
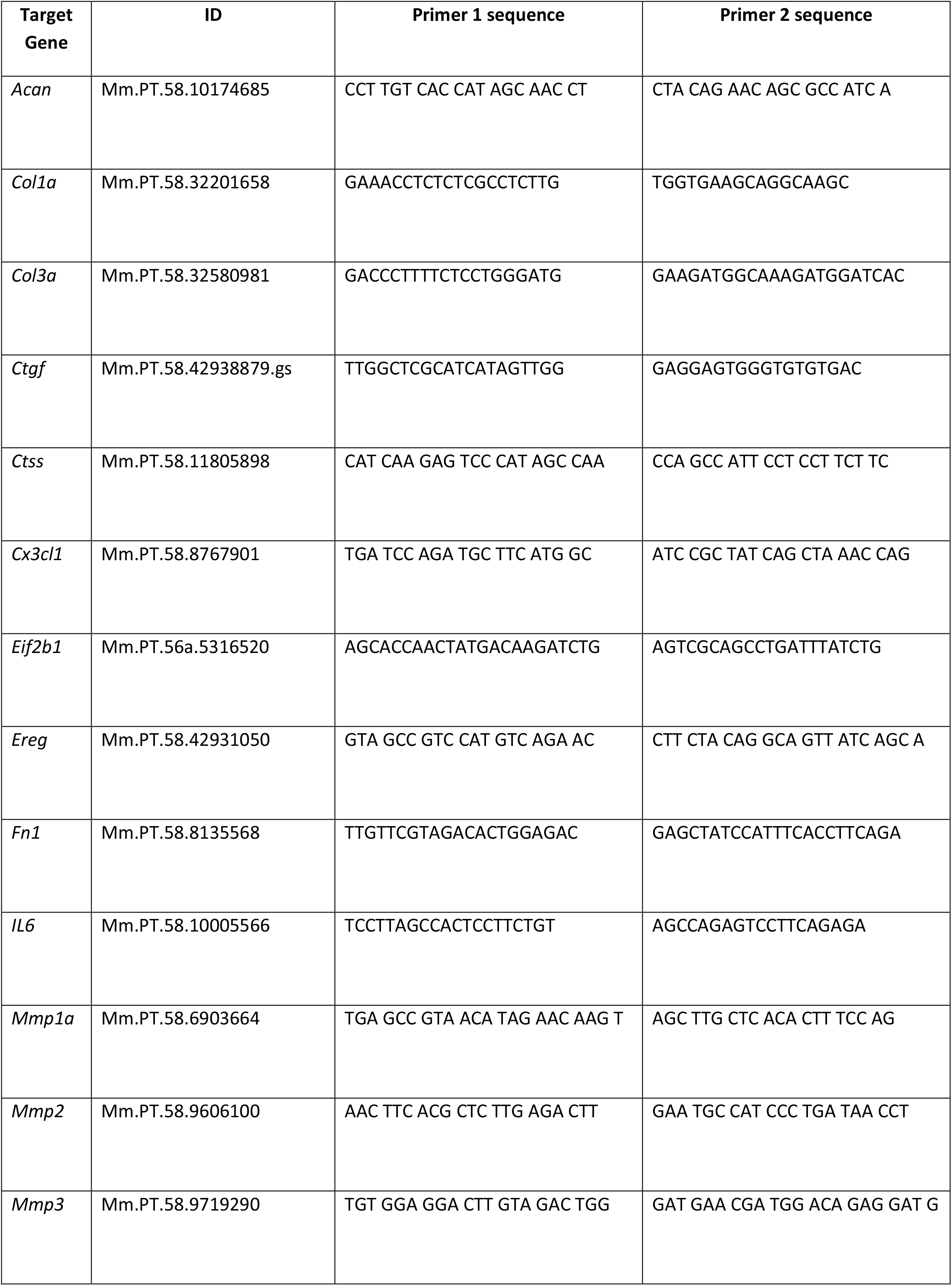
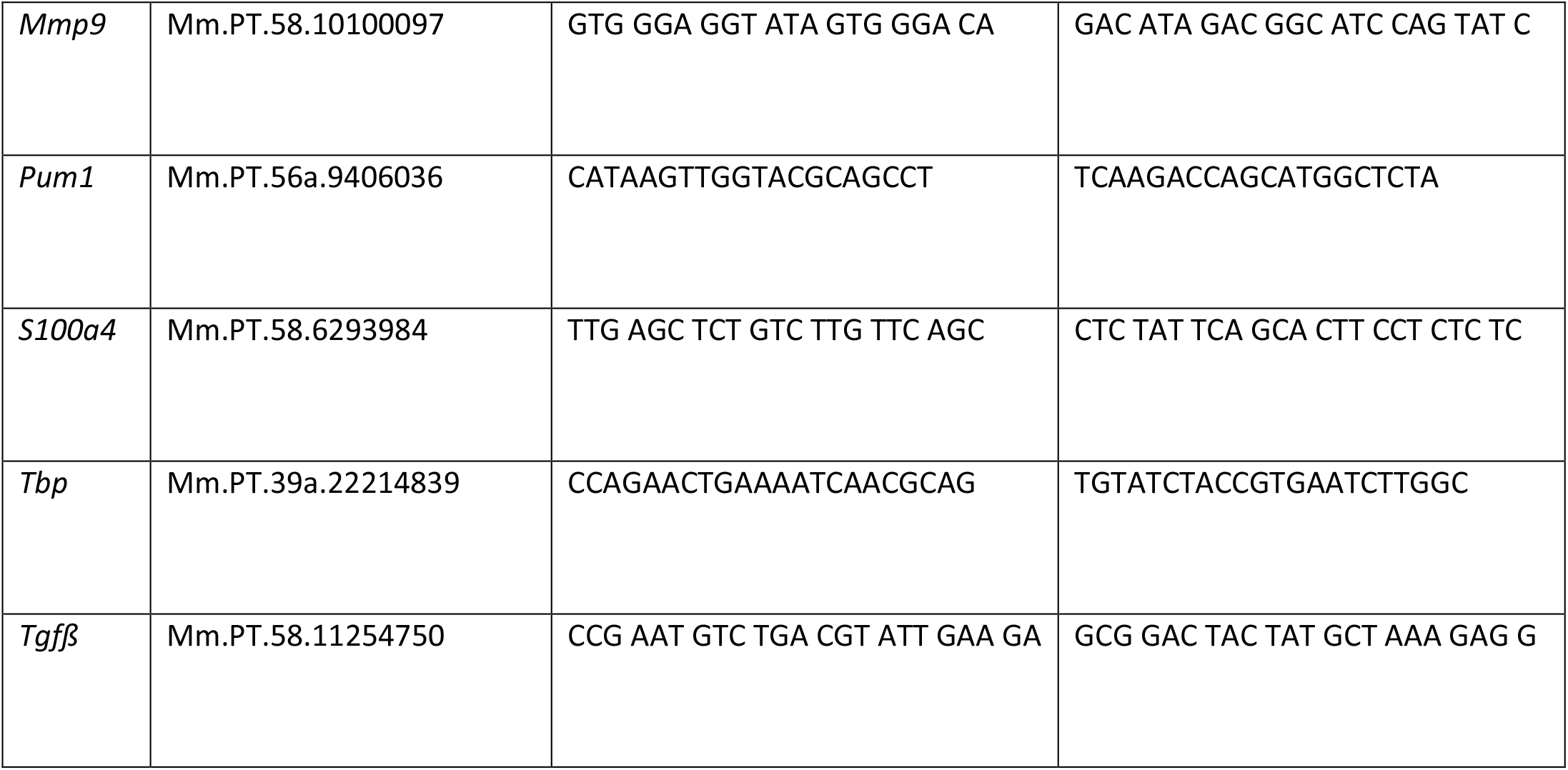
List of assays and respective primer sequences used for RT-qPCR analysis

#### Study approval

All animal experiments and procedures were carried out in accordance with Austrian laws on animal experimentation and were approved by Austrian regulatory authorities (Project License No. BMWFW-66.019/0023-WF/V/3b/2017). The Paracelsus 10.000 study was approved by the local Ethics committee P10-415-E/1521/3-2012. Written informed consent was obtained from all participants prior to inclusion in the study.

#### Paracelsus 10.000 study

As part of a large-scale epidemiological health study initiated and conducted by the University Clinic Salzburg and the Paracelsus Medical University Salzburg, 10.000 participants aged between 40 and 69 years were queried regarding their general health and life style habits. In addition to collecting basic physical parameters such as average blood pressure, weight, and height, several sociodemographic factors were surveyed. Two extensive questionnaires and an in-depth interview were used to obtain information on lifestyle, work-related physical activity (e.g. heavy, intermediate, and little physical loading of the shoulders), and to collect data on patient’s history of diseases, medical interventions and medication. All variables used in this analysis were obtained in one of three ways: either from a questionnaire filled in by the participants themselves, from a structured face-to-face interview or from direct physical measurement. Body mass index (BMI) was calculated from direct measurement of height and weight on the study site. We defined obesity to be present when BMI exceeded 30 kg/m^2.

In the structured interview the interviewer asked whether certain diseases had ever been diagnosed by a doctor. The ones used for the purpose of our study were allergic rhinitis, asthma bronchiale, diabetes type 1, diabetes type 2, chronic obstructive pulmonary disease (COPD), rheumatoid arthritis, psoriasis, colitis ulcerosa and Crohn’s disease. Since diabetes type 1 was rare, we pooled both types of diabetes together under „diabetes”. The same accounted for colitis ulcerosa and Crohn’s disease which we therefore subsumed under „inflamatory bowel diseases” (IBD).

In the questionnaire participants were asked whether they experienced any allergies or intolerances. Possible answers were „No”, „Yes, allergy to pollen”, „Yes, allergy to mites”, „Yes, allergy to animal hair”, „Yes, food allergy against nuts, fruits, fish or seafood”, „Yes, intolerance against Fructose, Lactose or Histamine” „Yes, allergy against insects”, and, „Yes, allergy against drugs”. It was possible to choose more than one answer. We defined a participant as suffering from allergy if either a confirmed diagnosis of allergic rhinitis or asthma bronchiale was reported or „Yes” to any of the possible allergies in the self-report was reported. Intolerance against fructose, lactose or histamine was however excluded from the analysis.

Use of medication was also assessed during the structured interview. Medication was classified into different categories and only use of glucocorticoids was included. We differentiated between systemic, inhalative and local use of glucocorticoids, with the later subsuming topical applications as well as injections. For the purposes of this analysis we defined glucocorticoid use as either systemic or inhalative use or injections, however excluding topical application. We further defined glucocorticoid use as current use only and did not consider past use.

Cigarette smoking was also assessed via self-report. Participants were asked whether they currently or ever smoked cigarettes. They were further asked about the time span of smoking and the average amount of cigarettes consumed during that time, allowing for the calculation of package years (py).

Participants were also queried regarding any manual labour placing significant strain on the shoulder. Possible answers were: „Overhead work (e.g carpenter or painter)”, „heavy physical work on the left side (e.g mason or plumber)”, „light physical work (e.g. surgeon, electrician, truck driver or bus driver)”, „no physical work (cubicle work etc)”. For our purposes we considered overhead work to be relevant. Further, shoulder pain was assessed. Possible answers were „on the left side”, „on the right side”, „no nocturnal shoulder pain”. We considered nocturnal shoulder pain to be present if one of the first two answers was given. Finally, participants were asked if they were able to lift a carton of milk over their head and whether they had an accident leading to a shoulder injury.

The self-report further provided a free form field to list all surgeries ever performed on the participants. For our definition of tendinopathy, we manually selected relevant tendinopathy-related keywords in this field which could be classified into the following 3 subcategories: Carpal tunnel syndrome, tendon calcification, and tenosynovitis. We defined tendinopathy as either having had any surgery on this list, OR being unable to lift the carton of milk OR suffering from nocturnal shoulder pain. Based on these data, we filtered patients suffering from one of the reported allergies and displaying one of the above mentioned tendinopathy-associated features

### Statistical analysis

We explored the influence of allergy on tendinopathy using logistic regression modelling with tendinopathy as the outcome. We fitted one univariate model with only allergy as the predictor (Model 1) and two multivariate models. In the first multivariate model we adjusted for obesity, age and sex (Model 2). In a third model we further added glucocorticoid use, cigarette smoking, the two physical risk factors of accidents with shoulder injury and overhead work as well as comorbidities that are either known risk factors for tendinopathy, and like allergy are also associated with chronic inflammation, or can also lead to a subscription of glucocorticoids. These were COPD, psoriasis, rheumatoid arthritis, and inflammatory bowel diseases. Table S1 provides information about the missingness of the variables used in Model 3 (see supplements).

In order to evaluate the data set as objectively as possible, we also performed sensitivity analyses. To this end, we changed the definition of the outcome. We selected four different outcome variables combining the above described tendinopathy related features in different ways: 1) nocturnal shoulder pain OR inability to lift a milk carton overhead (N=2303); 2) nocturnal shoulder pain AND inability to lift a milk carton overhead (N=426); 3) one tendinopathy-related keyword from the list OR nocturnal shoulder pain OR inability to lift a milk carton overhead (N=2355); 4) one tendinopathy-related keyword from the list OR nocturnal shoulder pain AND inability to lift a milk carton overhead (N=500). Since there was no significant difference between outcome variables 1 and 2 compared to outcomes using the variables 3 and 4, only the latter results are reported, with variable 3 being used for our primary analysis, and variable 4 being used in the supplements.

All statistical analyses of the Paracelsus 10.000 data set were conducted using R (Version 4.0.2).

For the statistical analyses of all other experiments GraphPad Prism v.5.04 (La Jolla, CA, USA) was used. Numerical data is presented as means ± SEM. One-way analysis of variance (ANOVA) for multiple comparisons and 2-sample t-test for pair-wise comparisons were employed after confirming normal distribution of the data (D’Agostino or Pearson omnibus normality test or Shapiro-Wilkinson test). Non-parametric statistics were utilized when the above assumption was violated and consequently Kruskal–Wallis test for multiple comparisons or Mann–Whitney test to determine two-tailed p-value samples was carried out. To analyse for differences in collagen fibril size distribution between groups the Linear mixed model fit by maximum likelihood (‘lmerMod’) was applied. Numbers of biological replicates and statistical tests applied are reported in the figure legends. Statistical significance was set at α = 0.05.

## Acknowledgements

We would like to thank Dr. Vanessa Frey for providing access to the data of the Paracelsus 10.000 study and Dr. Wolfgang Forstmeier for his assistance with statistics. We kindly acknowledge the support of the Fund for the Advancement of Scientific Research at Paracelsus Medical University (PMU-FFF E-15/22/115-LEK).

## Author contributions

CL, HT, and AT conceptualized the project, designed the experiments, and analysed the data. CL performed western blot and TEM analysis, and drafted the manuscript. GS and RG established 3D tendon-like constructs and performed real-time qPCR. HT performed biomechanical testing and immunofluorescence staining. AW performed cryosectionning and polarization microscopy. PL performed statistical analysis of the human data set. NW and DJ assisted with setup of the animal study and collection of tissue samples. BK, RW and LA established the allergy mouse model. RW performed blood cytokine analysis. ET, BI, and BP designed and conducted the human Paracelsus 10.000 study. AT supervised the project, reviewed and edited the manuscript.

## The paper explained

### Problem

The etiology of tendinopathies is poorly understood. A potential contribution of inflammation in onset and progression of the disease is still controversially debated. Risk factors associated with the development of tendinopathies such as diabetes, obesity or rheumatoid arthritis are known to go along with a systemic inflammation. Whether also the presence of an allergy and the accompanying systemic inflammation affects tendon properties has not been investigated so far.

### Results

Induction of an allergic response in mice led to a significant upregulation of circulating blood cytokines. Tendons of these animals displayed signs of reduced tissue quality, including a significant reduction of the elastic modulus and tensile stress, but also alterations of the tendon matrix and increased expression of matrix-remodelling and inflammation-associated markers. Importantly, a large health survey study revealed that the presence of an allergy increases the risk to develop a tendinopathy.

### Impact

So far allergic diseases have not been considered a risk factor for tendinopathies. Knowing that the presence of a systemic inflammation alone is sufficient to negatively impact on tendons adds to a better understanding of the patho-mechanisms of the disease.

## Expanded view figure legends

**Fig. EV1:**
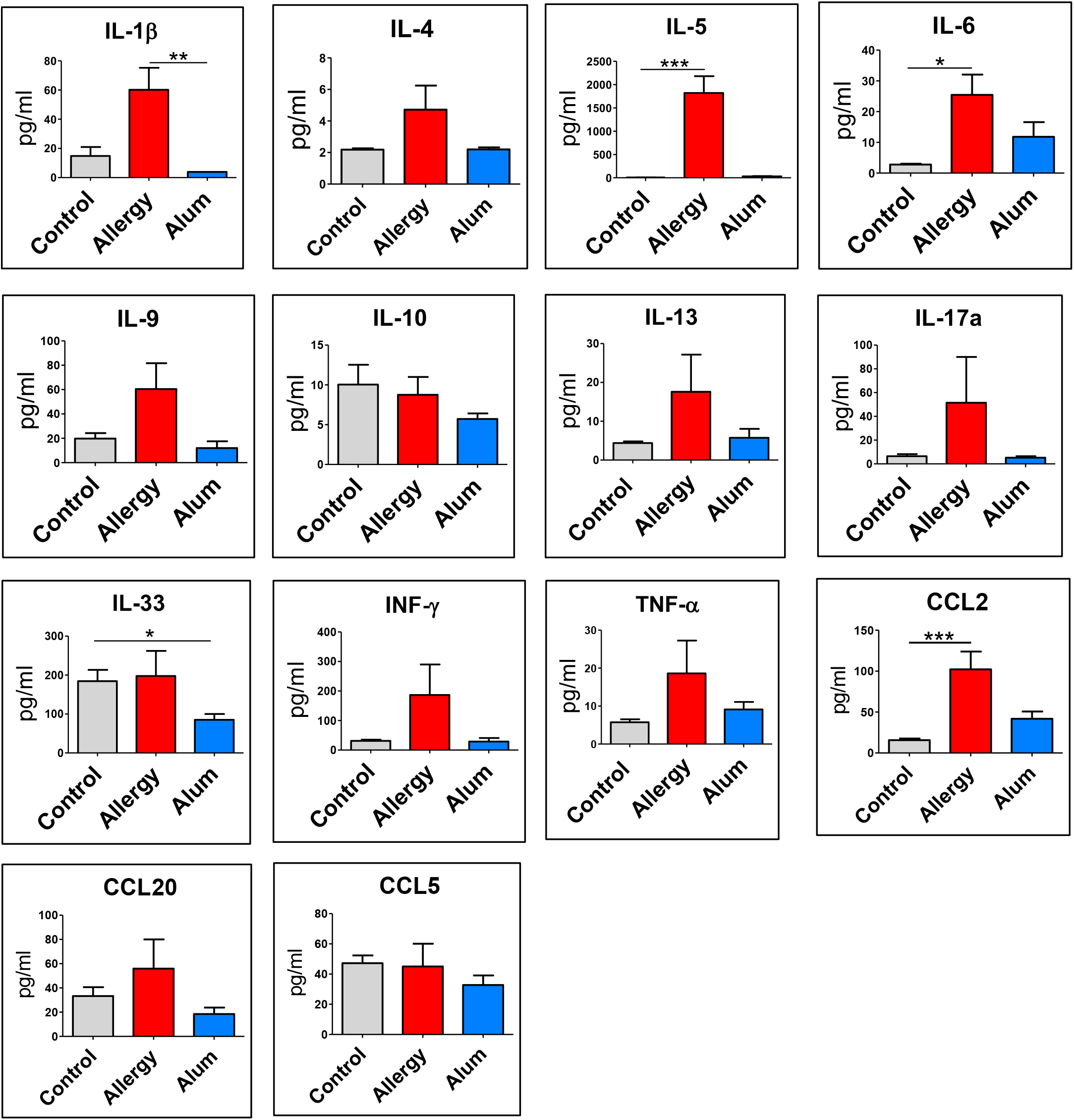
Selected serum cytokines/chemokines measured by a multiplex bead array 1 day after the sensitization phase. Data are shown as Mean + SEM, *p<0.05, **p<0.01, ***p<0.001, One-way ANOVA (Kruskal-Wallis and Dunn’s Multiple Comparison Test), n (control/allergy) = 16, n (Alum) = 6.

**Fig. EV2:**
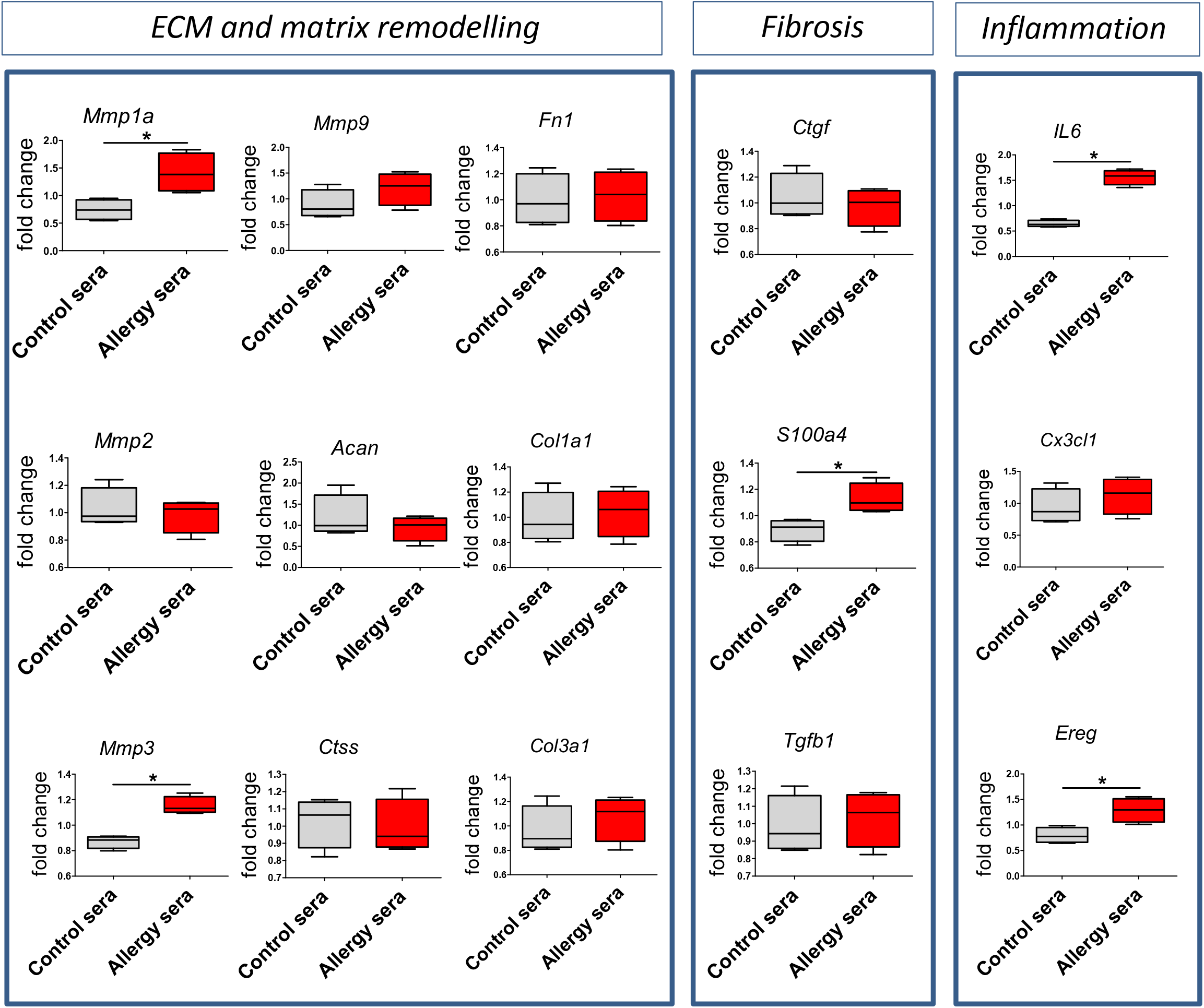
qPCR analysis of 3D tendon-like constructs. Treatment of tendon-like constructs with a pool of sera from allergic mice result in upregulation of selected target genes; *p<0.05, **p < 0.01, Mann-Whitney U test, n = 4. Bars represent mean ± SEM.

**Table EV1.**
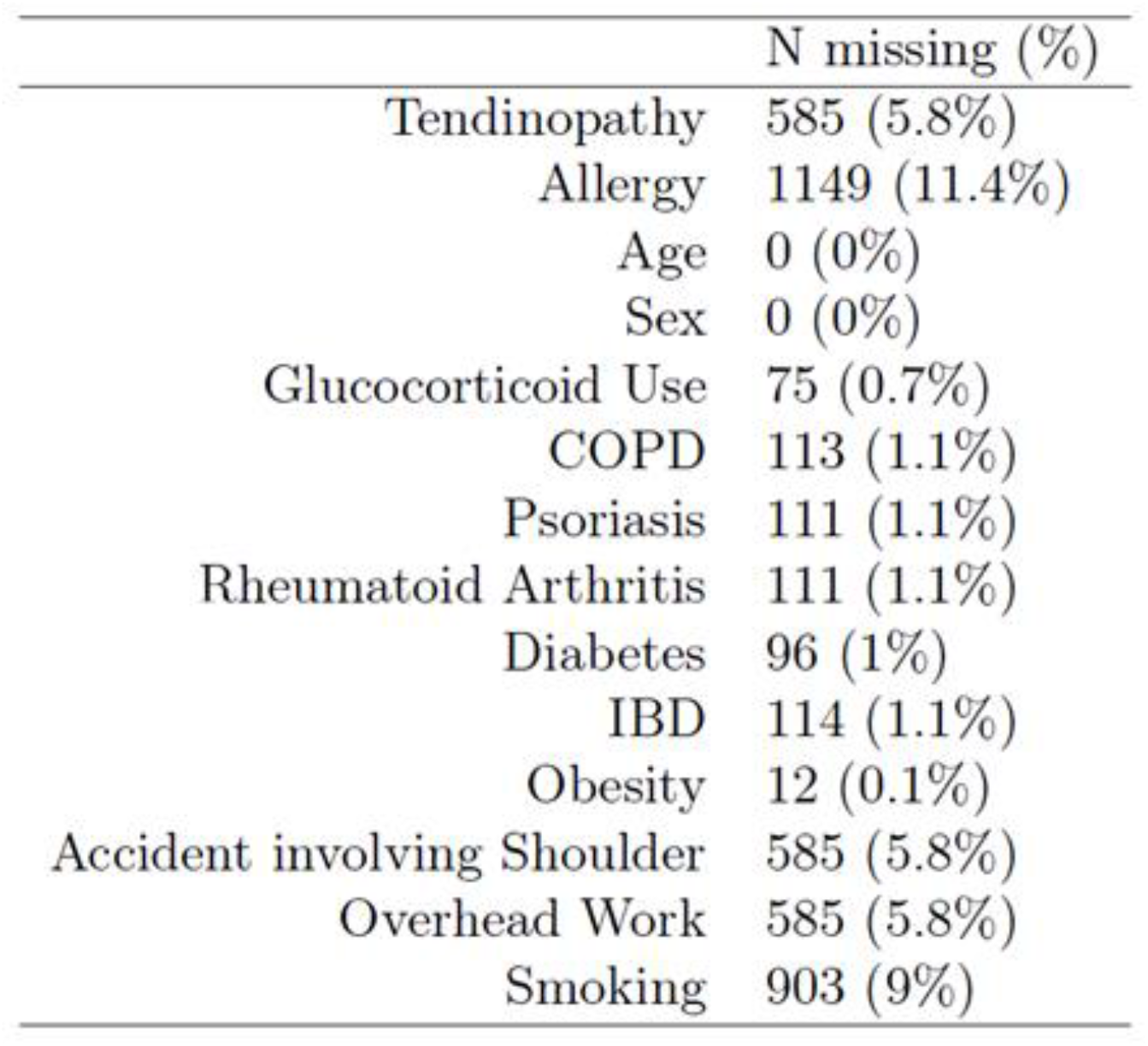
Number and percentages of missing observations for each variable. Table S1 provides information about the missingness of the variables used in Model 3. The 5.8% missing for tendinopathy, accidents involving the shoulder and overhead work are due to participants that did not fill in the self-report at all. The number of missing values for allergy is even higher, owing to the fact that the questions about allergy were only added to the questionnaire sometime after enrollment into the study began. The missingness rate for smoking is also larger than the 5.8% generally missing from the self-report. The excess 3.2% missing are due to information needed to calculate package years being insufficient or ambiguous.

**Table EV2.**
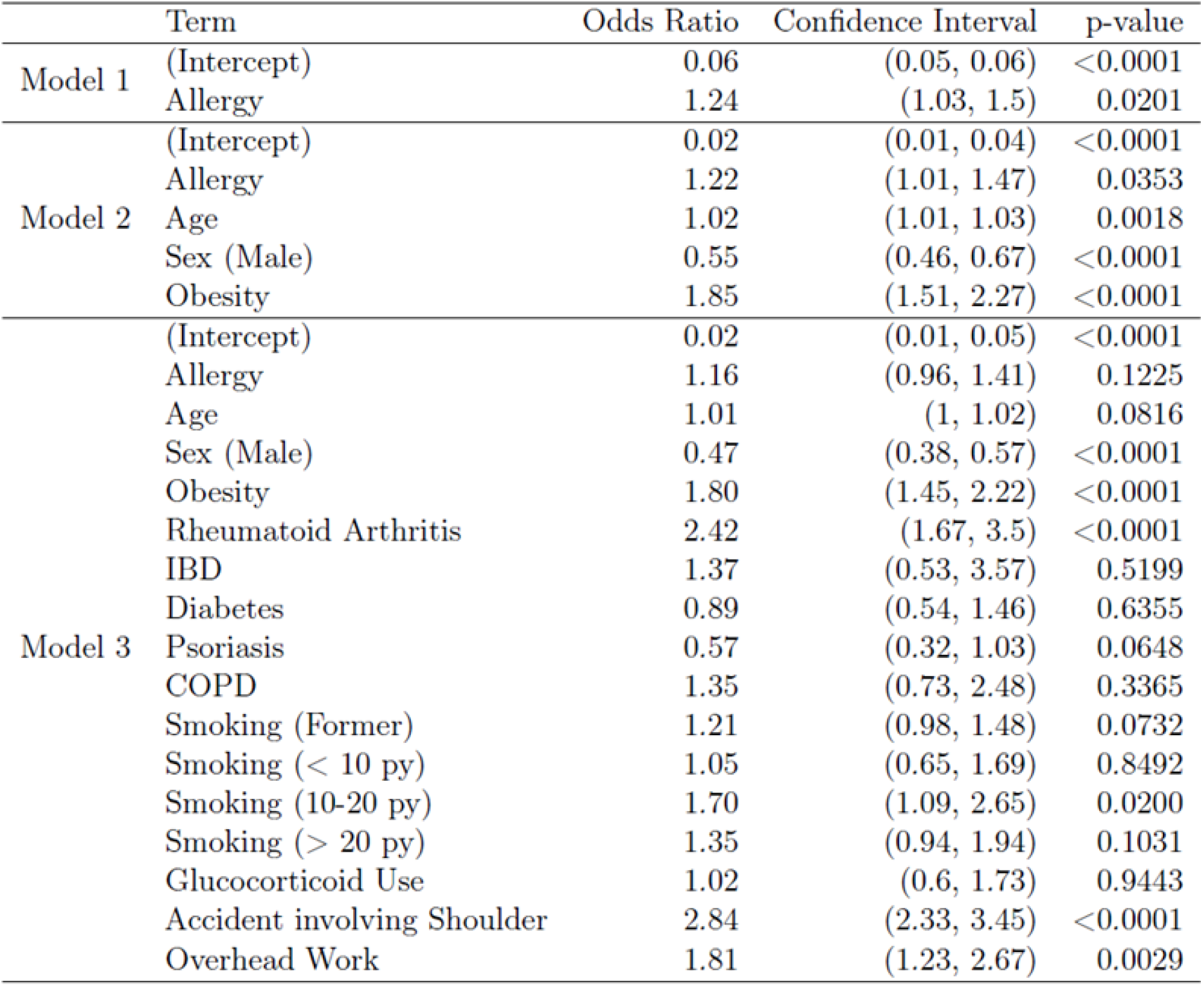
Odds Ratios, 95% Confidence Intervals for the Odds Ratio and p-values for all terms in the three models when the outcome is presence of a tendon surgery OR inability to lift a carton of milk overhead AND nocturnal shoulder pain.

|  | Term | Odds Ratio | Confidence Interval | p-value |
| --- | --- | --- | --- | --- |
| Model 1 | (Intercept) | 0.06 | (0.05, 0.06) | <0.0001 |
|  | Allergy | 1.24 | (1.03, 1.5) | 0.0201 |
| Model 2 | (Intercept) | 0.02 | (0.01, 0.04) | <0.0001 |
|  | Allergy | 1.22 | (1.01, 1.47) | 0.0353 |
|  | Age | 1.02 | (1.01, 1.03) | 0.0018 |
|  | Sex (Male) | 0.55 | (0.46, 0.67) | <0.0001 |
|  | Obesity | 1.85 | (1.51, 2.27) | <0.0001 |
| Model 3 | (Intercept) | 0.02 | (0.01, 0.05) | <0.0001 |
|  | Allergy | 1.16 | (0.96, 1.41) | 0.1225 |
|  | Age | 1.01 | (1, 1.02) | 0.0816 |
|  | Sex (Male) | 0.47 | (0.38, 0.57) | <0.0001 |
|  | Obesity | 1.80 | (1.45, 2.22) | <0.0001 |
|  | Rheumatoid Arthritis | 2.42 | (1.67, 3.5) | <0.0001 |
|  | IBD | 1.37 | (0.53, 3.57) | 0.5199 |
|  | Diabetes | 0.89 | (0.54, 1.46) | 0.6355 |
|  | Psoriasis | 0.57 | (0.32, 1.03) | 0.0648 |
|  | COPD | 1.35 | (0.73, 2.48) | 0.3365 |
|  | Smoking (Former) | 1.21 | (0.98, 1.48) | 0.0732 |
|  | Smoking (< 10 py) | 1.05 | (0.65, 1.69) | 0.8492 |
|  | Smoking (10-20 py) | 1.70 | (1.09, 2.65) | 0.0200 |
|  | Smoking (> 20 py) | 1.35 | (0.94, 1.94) | 0.1031 |
|  | Glucocorticoid Use | 1.02 | (0.6, 1.73) | 0.9443 |
|  | Accident involving Shoulder | 2.84 | (2.33, 3.45) | <0.0001 |
|  | Overhead Work | 1.81 | (1.23, 2.67) | 0.0029 |

**Table EV3.** List of assays and respective primer sequences used for RT-qPCR analysis

| Cytokine | Company | Catalog number |
| --- | --- | --- |
| Recombinant rat IL1B | Peprotech | 400-01B |
| Recombinant rat IL4 | Peprotech | 400-04 |
| Recombinant rat IL5 | Peprotech | 400-05 |
| Recombinant rat IL6 | Peprotech | 400-06 |
| Recombinant murine IL17A | Peprotech | 210-17 |
| Recombinant rat INFγ | Peprotech | 400-20 |
| Recombinant rat TNFα | Peprotech | 400-14 |

## References

1. Hopkins, C., et al., Critical review on the socio-economic impact of tendinopathy. Asia Pac J Sports Med Arthrosc Rehabil Technol, 2016. 4: p. 9–20.

2. Thomopoulos, S., et al., Mechanisms of tendon injury and repair. Journal of orthopaedic research: official publication of the Orthopaedic Research Society, 2015. 33(6): p. 832–9.

3. Cho, N.S., et al., Retear patterns after arthroscopic rotator cuff repair: single-row versus suture bridge technique. Am J Sports Med, 2010. 38(4): p. 664–71.

4. Choi, S., et al., Factors associated with clinical and structural outcomes after arthroscopic rotator cuff repair with a suture bridge technique in medium, large, and massive tears. J Shoulder Elbow Surg, 2014. 23(11): p. 1675–81.

5. Kirchgesner, T., et al., Drug-induced tendinopathy: from physiology to clinical applications. Joint Bone Spine, 2014. 81(6): p. 485–92.

6. Dean, B.J., et al., Are inflammatory cells increased in painful human tendinopathy? A systematic review. Br J Sports Med, 2016. 50(4): p. 216–20.

7. Millar, N.L., et al., Inflammation is present in early human tendinopathy. Am J Sports Med, 2010. 38(10): p. 2085–91.

8. Rees, J.D., M. Stride, and A. Scott, Tendons--time to revisit inflammation. Br J Sports Med, 2014. 48(21): p. 1553–7.

9. Speed, C., Inflammation in Tendon Disorders. Adv Exp Med Biol, 2016. 920: p. 209–20.

10. Franceschi, F., et al., Obesity as a risk factor for tendinopathy: a systematic review. Int J Endocrinol, 2014. 2014: p. 670262.

11. Gaida, J.E., et al., Is adiposity an under-recognized risk factor for tendinopathy? A systematic review. Arthritis Rheum, 2009. 61(6): p. 840–9.

12. Beason, D.P., et al., Cumulative effects of hypercholesterolemia on tendon biomechanics in a mouse model. Journal of orthopaedic research: official publication of the Orthopaedic Research Society, 2011. 29(3): p. 380–3.

13. Titchener, A.G., et al., Comorbidities in rotator cuff disease: a case-control study. J Shoulder Elbow Surg, 2014. 23(9): p. 1282–8.

14. Ranger, T.A., et al., Is there an association between tendinopathy and diabetes mellitus? A systematic review with meta-analysis. Br J Sports Med, 2016. 50(16): p. 982–9.

15. Ribbans, W.J. and P.D. Angus, Simultaneous bilateral rupture of the quadriceps tendon. Br J Clin Pract, 1989. 43(3): p. 122–5.

16. Dungey, M., et al., Inflammatory factors and exercise in chronic kidney disease. Int J Endocrinol, 2013. 2013: p. 569831.

17. Battery, L. and N. Maffulli, Inflammation in overuse tendon injuries. Sports Med Arthrosc, 2011. 19(3): p. 213–7.

18. Petersen, A.M. and B.K. Pedersen, The anti-inflammatory effect of exercise. J Appl Physiol (1985), 2005. 98(4): p. 1154–62.

19. Pistone, G., et al., Achilles tendon ultrasonography may detect early features of psoriatic arthropathy in patients with cutaneous psoriasis. Br J Dermatol, 2014. 171(5): p. 1220–2.

20. Frizziero, A., et al., Foot tendinopathies in rheumatic diseases: etiopathogenesis, clinical manifestations and therapeutic options. Clin Rheumatol, 2013. 32(5): p. 547–55.

21. Kaur, S., et al., Comparative study of systemic inflammatory responses in psoriasis vulgaris and mild to moderate allergic contact dermatitis. Dermatology, 2012. 225(1): p. 54–61.

22. Lai, N.S., et al., Association of rheumatoid arthritis with allergic diseases: A nationwide population-based cohort study. Allergy Asthma Proc, 2015. 36(5): p. 99–103.

23. Prescott, S.L., Early-life environmental determinants of allergic diseases and the wider pandemic of inflammatory noncommunicable diseases. J Allergy Clin Immunol, 2013. 131(1): p. 23–30.

24. Klein, B., et al., Allergy Enhances Neurogenesis and Modulates Microglial Activation in the Hippocampus. Front Cell Neurosci, 2016. 10: p. 169.

25. Lehner, C., et al., Tenophages: a novel macrophage-like tendon cell population expressing CX3CL1 and CX3CR1. Dis Model Mech, 2019.

26. Lui, P.P.Y., Tendinopathy in diabetes mellitus patients-Epidemiology, pathogenesis, and management. Scand J Med Sci Sports, 2017. 27(8): p. 776–787.

27. Macchi, M., et al., Obesity Increases the Risk of Tendinopathy, Tendon Tear and Rupture, and Postoperative Complications: A Systematic Review of Clinical Studies. Clin Orthop Relat Res, 2020.

28. Hahn, C., et al., Inhibition of the IL-4/IL-13 receptor system prevents allergic sensitization without affecting established allergy in a mouse model for allergic asthma. J Allergy Clin Immunol, 2003. 111(6): p. 1361–9.

29. Hirose, K., et al., Allergic airway inflammation: key players beyond the Th2 cell pathway. Immunol Rev, 2017. 278(1): p. 145–161.

30. Wang, Y.H. and Y.J. Liu, The IL-17 cytokine family and their role in allergic inflammation. Curr Opin Immunol, 2008. 20(6): p. 697–702.

31. Kordulewska, N.K., et al., Cytokines concentrations in serum samples from allergic children-Multiple analysis to define biomarkers for better diagnosis of allergic inflammatory process. Immunobiology, 2018. 223(11): p. 648–657.

32. Hernandez, P., et al., Severe Burn-Induced Inflammation and Remodeling of Achilles Tendon in a Rat Model. Shock, 2018. 50(3): p. 346–350.

33. Choi, R.K., et al., Chondroitin sulphate glycosaminoglycans contribute to widespread inferior biomechanics in tendon after focal injury. J Biomech, 2016. 49(13): p. 2694–2701.

34. Samiric, T., et al., Changes in the composition of the extracellular matrix in patellar tendinopathy. Matrix Biol, 2009. 28(4): p. 230–6.

35. Fu, S.C., K.M. Chan, and C.G. Rolf, Increased deposition of sulfated glycosaminoglycans in human patellar tendinopathy. Clin J Sport Med, 2007. 17(2): p. 129–34.

36. Lehner, C., et al., Tenophages: a novel macrophage-like tendon cell population expressing CX3CL1 and CX3CR1. Dis Model Mech, 2019. 12(12).

37. Fu, S.C., et al., Increased expression of matrix metalloproteinase 1 (MMP1) in 11 patients with patellar tendinosis. Acta Orthop Scand, 2002. 73(6): p. 658–62.

38. Alfredson, H., et al., cDNA-arrays and real-time quantitative PCR techniques in the investigation of chronic Achilles tendinosis. J Orthop Res, 2003. 21(6): p. 970–5.

39. Sejersen, M.H., et al., Proteomics perspectives in rotator cuff research: a systematic review of gene expression and protein composition in human tendinopathy. PLoS One, 2015. 10(4): p. e0119974.

40. Etzerodt, A. and S.K. Moestrup, CD163 and inflammation: biological, diagnostic, and therapeutic aspects. Antioxid Redox Signal, 2013. 18(17): p. 2352–63.

41. Skytthe, M.K., J.H. Graversen, and S.K. Moestrup, Targeting of CD163(+) Macrophages in Inflammatory and Malignant Diseases. Int J Mol Sci, 2020. 21(15).

42. Lam, D., S. Lively, and L.C. Schlichter, Responses of rat and mouse primary microglia to pro- and anti-inflammatory stimuli: molecular profiles, K(+) channels and migration. J Neuroinflammation, 2017. 14(1): p. 166.

43. Londos, C., et al., Perilipins, ADRP, and other proteins that associate with intracellular neutral lipid droplets in animal cells. Semin Cell Dev Biol, 1999. 10(1): p. 51–8.

44. Rosas-Ballina, M., et al., Classical Activation of Macrophages Leads to Lipid Droplet Formation Without de novo Fatty Acid Synthesis. Front Immunol, 2020. 11: p. 131.

45. den Brok, M.H., et al., Lipid Droplets as Immune Modulators in Myeloid Cells. Trends Immunol, 2018. 39(5): p. 380–392.

46. D’Addona, A., et al., Inflammation in tendinopathy. Surgeon, 2017. 15(5): p. 297–302.

47. Leong, H.T., et al., Risk factors for rotator cuff tendinopathy: A systematic review and meta-analysis. J Rehabil Med, 2019. 51(9): p. 627–637.

48. Rechardt, M., et al., Lifestyle and metabolic factors in relation to shoulder pain and rotator cuff tendinitis: a population-based study. BMC Musculoskelet Disord, 2010. 11: p. 165.

49. Wendelboe, A.M., et al., Associations between body-mass index and surgery for rotator cuff tendinitis. J Bone Joint Surg Am, 2004. 86(4): p. 743–7.

50. Gehwolf, R., et al., 3D-Embedded Cell Cultures to Study Tendon Biology. Methods Mol Biol, 2019.

51. Lehner, C., et al., The blood-tendon barrier: identification and characterisation of a novel tissue barrier in tendon blood vessels. Eur Cell Mater, 2016. 31: p. 296–311.

52. Vandesompele, J., et al., Accurate normalization of real-time quantitative RT-PCR data by geometric averaging of multiple internal control genes. Genome Biol, 2002. 3(7): p. Research0034.

